# Systematic mapping of chimera-permissive sites by CRISPR-guided PAM scanning

**DOI:** 10.64898/2026.05.06.723292

**Authors:** Jacinda Pujols, Kyutae D. Lee, Nick J. Kapolka, Sam Taylor, Bruno Colon, Daniel G. Isom

## Abstract

Engineered protein chimeras enable new biological functions but remain difficult to design due to context-dependent constraints on insertion tolerance and the need to preserve host protein function. Here, we report CRISPR-guided protospacer adjacent motif (PAM) scanning in yeast to map chimera-permissive sites in living cells. We apply this approach to peptide and reporter insertions. In the first application, we generated 91 insertion chimeras encoding a defined protease cleavage sequence across six components of a model G protein–coupled receptor (GPCR) signaling pathway. Sixty-three percent of sites retain signaling, identifying positions that preserve host function and reveal broad, position-dependent tolerance. Coupling insertional scanning with cognate proteases enables site-resolved mapping of in-cell accessibility, distinguishing protected and exposed regions and defining EX1- and EX2-like regimes. These chimeras are responsive to proteolysis and pharmacological inhibition, enabling reversible control of protein activity. In a second application, we scanned 32 positions in yeast Ste2 and human A2A and MTNR1A receptors to engineer bi-functional chimeras that retain native function while incorporating reporter activity. Together, these results establish PAM scanning as a scalable, protein-agnostic framework for mapping insertion tolerance, interrogating protein accessibility *in vivo*, and enabling scalable ground-truth benchmarking of predictive chimera engineering.

## Introduction

Protein chimeras are single polypeptides assembled from domains or sequence elements derived from distinct proteins. In nature, exon shuffling, gene fusion, recombination, and chromosomal rearrangement can recombine pre-existing modules to generate new regulatory, catalytic, and signaling functions, thereby accelerating evolutionary innovation. The same processes can also drive disease. Recurrent oncogenic fusion proteins such as BCR-ABL1, PML-RARA, and EML4-ALK rewire signaling or transcriptional control to promote malignant transformation^1^. Natural chimeras, therefore, illustrate both the creative and pathological consequences of domain recombination, a principle increasingly exploited through protein engineering.

Engineered protein chimeras have become indispensable tools for converting otherwise inaccessible molecular and cellular events into measurable signals. Fusion-based designs, including early GFP and luciferase reporters, transformed cell biology by enabling direct visualization of protein localization, abundance, and dynamics in living systems^2–4^. Insertion-based designs extended this principle, yielding biosensors such as GCaMP and reporters for voltage, cAMP, and kinase activity^5–7^. Chimeric G protein–coupled receptor (GPCR) sensors, generated by inserting fluorescent domains into intracellular loops, further enabled real-time measurement of ligand binding, receptor activation, and effector recruitment^8–14^. Together, these studies established that strategically placed insertions can serve as quantifiable readouts of changes in protein conformation, function, or location.

The modularity of protein chimeras has also enabled major therapeutic advances. Approved biologics such as etanercept, aflibercept, and abatacept combine receptor or regulatory domains with immunoglobulin Fc regions to improve stability, half-life, and efficacy^15,16^. Chimeric monoclonal antibodies, such as rituximab, similarly exploit modular architectures to combine antigen specificity with immune effector function and have transformed the treatment of lymphoma and autoimmune diseases^17,18^. More recently, chimeric antigen receptors used in CAR-T therapies redirect immune cells against cancer, with transformative clinical benefits^19–21^. These examples underscore the therapeutic power of recombining protein modules with defined functions.

Despite their utility, functional protein chimeras remain challenging to design because protein engineering is highly context dependent. Terminal fusion proteins are often more tractable, as appended domains may spare the host fold. By contrast, insertion chimeras, in which one domain is embedded within another, face stricter constraints to preserve folding, allostery, and native activity while enabling productive interdomain communication. Permissive insertion sites are often sparse and remain difficult to predict solely from sequence, structure, or computational models. Therefore, defining such sites remains a bottleneck in biosensor engineering, synthetic biology, and therapeutic protein design.

Here, we develop a systematic framework for mapping chimera-permissive sites using Clustered Regularly Interspaced Short Palindromic Repeats (CRISPR)-guided protospacer adjacent motif (PAM) scanning. By leveraging programmable genome engineering to create dense insertion opportunities across coding sequences, we identify positions that tolerate domain integration in living cells across diverse protein classes and signaling systems. We apply this approach to define chimeric tolerance *in vivo* across a G protein-coupled receptor (GPCR) signaling pathway, map the in-cell biophysical accessibility of peptide insertions, and identify permissive sites in human GPCRs for biosensor-like readouts. More broadly, this framework enables large-scale functional interrogation of insertional tolerance, reveals principles governing permissive protein architectures, and provides a generalizable approach to accelerate the design of next-generation chimeric proteins.

## Results

### Leveraging high-throughput CRISPR in yeast to explore chimera space

To systematically map chimera-permissive insertion sites in living cells, we sought a platform that combined precise genome editing with scalable functional screening. CRISPR enables targeted sequence diversification and efficient editing at endogenous loci^22–25^. These advantages are especially pronounced in *Saccharomyces cerevisiae* (hereafter yeast), where robust homologous recombination, facile genetics, and scalability enable high-throughput engineering, and where strains can be extensively humanized to reconstitute mammalian signaling pathways and interrogate human proteins^26–31^. We therefore developed a yeast-based CRISPR platform to define insertional tolerance at scale and establish chimeric, genetically encodable protein tools translatable to human, mouse, and other experimentally relevant model systems.

CRISPR targeting relies on a Cas9 nuclease, directed by a single-guide RNA, to create a predictable double-strand break in complementary genomic DNA adjacent to a PAM site (Fig. 1a). Because PAM sites are distributed throughout coding sequences, they provide a programmable set of genomic entry points for systematic insertional scanning. We leveraged this constraint as a feature, using PAM availability to nominate codon-resolved positions for domain integration across target genes.

**Fig. 1.**
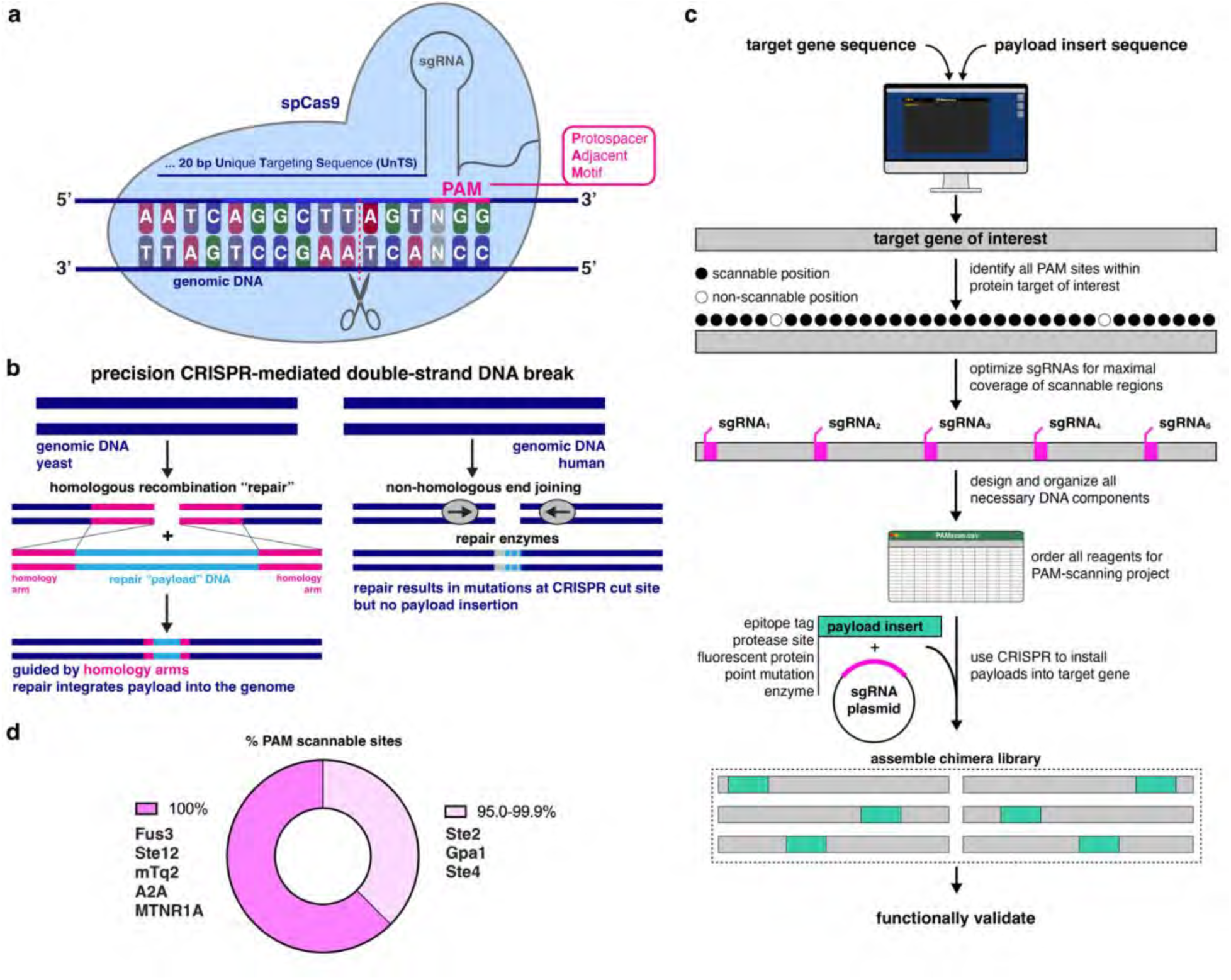
Principles and workflow of CRISPR-directed PAM-scanning. **a**, SpCas9 recognizes target DNA through a complementary 20-bp single guide RNA (sgRNA) and a protospacer adjacent motif (PAM; 5′-NGG-3′), with double-strand cleavage occurring three base pairs upstream of the PAM. **b**, Schematic of repair outcomes following CRISPR-induced double-strand breaks in yeast and human cells. In yeast (left), breaks are repaired by homologous recombination using a donor template with homology arms (pink), enabling precise integration of payload DNA (dark blue). In human cells (right), repair proceeds predominantly through non-homologous end joining, typically without payload incorporation. **c**, Overview of the PAM-scanning workflow. Target and payload sequences are input into a computational pipeline that identifies PAM sites, maps scannable positions, and optimizes sgRNAs to maximize coverage. Off-target potential is assessed by BLAST, and selected sgRNAs are assembled with donor templates to generate a complete design file. CRISPR-directed insertion produces libraries of chimeric variants, which are screened for functional outcomes. **d**, Fraction of PAM-scannable sites within each protein analyzed. Proteins include yeast mating pathway components (Ste2, Gpa1, Ste4, Fus3, Ste12), the reporter mTq2, and human GPCRs A2A and MTNR1A.

This strategy is particularly effective in yeast, which efficiently repairs CRISPR-induced breaks through homologous recombination using exogenous donor templates containing payload sequences flanked by homology arms (Fig. 1b). In contrast to mammalian systems, where non-homologous end joining frequently predominates, yeast supports precise and parallel installation of designed inserts at high efficiency^26–32^. We therefore implemented a PAM-scanning workflow that computationally identifies accessible PAM sites and designs guide RNAs and donor templates to generate libraries of insertion chimeras for functional analysis (Fig. 1c).

A key question is whether PAM sequence constraints leave enough accessible positions within coding sequences for meaningful insertional scanning. To address this, we quantified PAM-accessible codons across proteins examined in this study, including yeast signaling components, fluorescent reporters, and human GPCRs (Fig. 1d). We find that PAM sites occur frequently enough to enable broad, and often near-comprehensive, exploration of payload insertion opportunities. Under the editing rules used here, most proteins contained sufficient PAM-proximal positions to support systematic mapping of insertional tolerance rather than sparse interrogation of isolated sites.

### Automated design of PAM-defined insertion libraries

To enable systematic mapping of chimera-permissive sites, we developed software for automated identification of CRISPR-addressable insertion sites and the design of corresponding chimeric payloads. The workflow accepts a protein-coding open reading frame along with its flanking untranslated regions. For higher eukaryotic genes, which are frequently interrupted by multiple introns, reducing genomic structure to the spliced open reading frame substantially simplifies scanning and codon-resolved payload design while preserving the encoded protein sequence.

The pipeline then surveys both DNA strands for SpCas9 recognition motifs and identifies all candidate PAM sites. Because Cas9 cleavage can support editing across a local genomic window, each PAM site may access multiple nearby codons, allowing a single cut site to interrogate several potential insertion positions. This establishes a complete editable landscape for each target protein and enables either exhaustive library generation or structure-guided prioritization of informative sites. Here, we used experimentally determined protein structures and AlphaFold2 models to focus our codon sampling while retaining broad coverage across each protein.

For each target position, the software automatically generates the oligonucleotides required to construct guide RNA plasmids and donor templates. For small payloads, the insert sequence can be encoded directly within the primer overhangs. For larger payloads, primers are designed to amplify and assemble pre-existing inserts. This automation converts protein sequences into synthesis-ready reagent sets for arrayed PAM-scanning experiments.

As outlined in Fig. 1c, the algorithm further incorporates sequence safeguards to preserve desired edits after integration. Where possible, synonymous substitutions are introduced to eliminate the PAM while maintaining the encoded amino acid sequence, thereby preventing recurrent Cas9 cleavage after payload installation. When direct PAM disruption is not feasible, synonymous mismatches are introduced into the protospacer to reduce guide recognition while preserving host protein coding potential.

Candidate guides are then filtered by local BLAST+ analysis to minimize potential off-target alignments in the yeast genome. Remaining positions are ranked according to the distance between the Cas9 cut site and the codon of interest, and flanking primers are generated with universal suffixes compatible with our integration vectors. The final output is a genome-indexed catalog of PAM-scannable codons and corresponding reagents, enabling systematic installation of payloads across all target proteins. Across proteins examined here, the fraction of PAM-accessible codons was consistently high, supporting broad exploration of insertional tolerance (Fig. 1d).

### Global mapping of peptide insertion tolerance across the yeast mating pathway

As an initial application of PAM scanning, we systematically explored chimera tolerance and in-cell biophysical protection across a signaling pathway by introducing peptide inserts that confer susceptibility to a cognate protease. We implemented this strategy in the yeast mating pathway (Fig. 2a), an insulated GPCR signaling cascade that provides a quantitative readout for protein engineering^26–31^. In this system, the GPCR Ste2 detects extracellular mating pheromone (α-factor) and activates a heterotrimeric G protein complex comprising Gα (Gpa1) and Gβγ (Ste4/Ste18) subunits, with downstream signaling through the MAP kinase Fus3 and transcription factor Ste12 driving expression of the fluorescent reporter mTq2. Using the Hepatitis C virus (HCV) NS3/4A serine protease recognition sequence EDVVCCSMSY as a defined payload, we introduced peptide insertions into pathway components and assessed chimera tolerance via α-factor-stimulated mTq2 output.

**Fig. 2.**
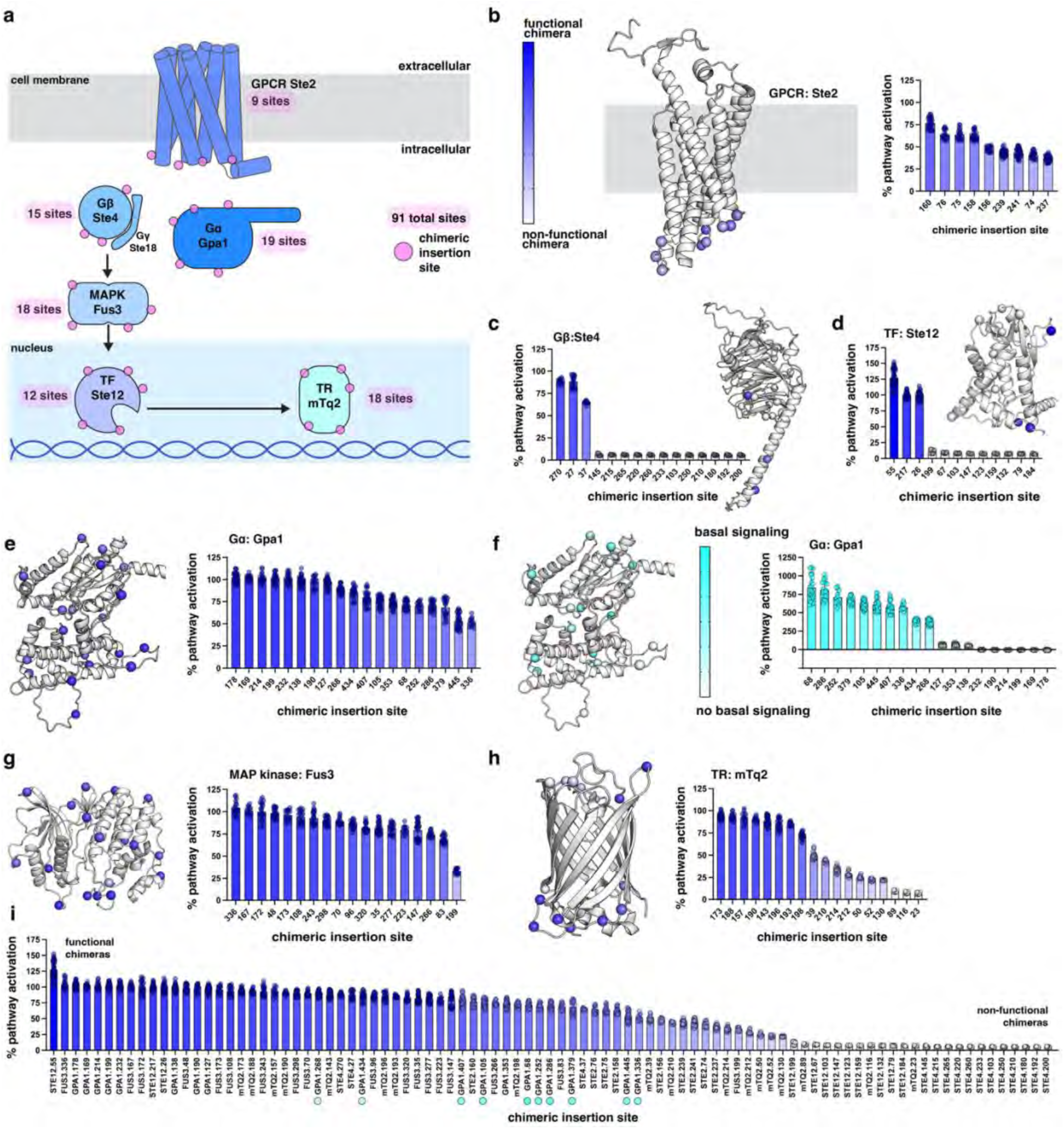
Global mapping of peptide insertion tolerance across the yeast mating pathway. **a**, Schematic of the yeast mating pathway indicating the number of PAM-scanned insertion sites in each signaling component. Chimeras incorporate the HCV NS3/4A protease substrate (EDVVCCSMSY). **b-e**,**g**,**h**, Signaling output of chimeras containing the 10-mer HCV protease substrate. Spheres (structural models) and bars (graphs) are color-coded from dark blue (functional) to white (non-functional). Pathway activation was induced with 10 μM α-factor (18–24 h) and quantified by endpoint fluorescence of the mTq2 transcriptional reporter. **f**, Basal signaling activity of Gpa1 (Gα) chimeras measured in the absence of α- factor (18–24 h). Spheres and bars are color-coded from bright cyan (high basal activity) to white (none). **i**, Waterfall plot summarizing signaling across all 91 insertion sites. Bar colors follow the key in panel b; non-tolerant Gpa1 chimeras with elevated basal activity are indicated by cyan circles. Data represent mean ± s.d. of ≥3 independent biological replicates, each with four technical replicates. See Supplementary Figs. S1, S2, S4, S5, S7–10, and S12–15 for additional data.

To achieve pathway-wide coverage, we generated 91 peptide insertion chimeras across the six pathway components (Fig. 2). We selected insertion sites from the full PAM-accessible landscape by integrating experimental structures with AlphaFold2 predictions, prioritizing positions distributed across each protein fold with a bias toward solvent-exposed loops and, for transmembrane receptors, intracellular regions outside the plasma membrane. Within the G protein module, we excluded Ste18, the Gγ subunit, due to its small size and obligate scaffolding role in stabilizing the Ste4 β-propeller. We instead treated the heterotrimer as a single structural unit while avoiding protein-protein interfaces.

We then assessed insertion tolerance across the pathway. Of the 91 sites, 57 (63%) retained measurable signaling above baseline, indicating broad tolerance to peptide insertion. All sites in Ste2 and Fus3 supported robust pathway signaling (Fig. 2b,g), yet signaling output spanned a continuum across tolerated sites, indicating that insertion position modulates pathway activity rather than simply permitting or abolishing function. Similarly, nearly all sites in the mTq2 transcriptional reporter (15 of 18) were tolerated (Fig. 2h).

Of the remaining 34 non-tolerated sites, 24 were mapped exclusively to Ste4, Ste12, and the mTq2 reporter (Fig. 2c,d,h). Disruption of the β-propeller fold or destabilization of the heterotrimeric complex in Ste4 chimeras led to 12 non-tolerated sites, suggesting reduced permissivity of this fold (Fig. 2c). Ste12 insertion positions were guided by a lower-confidence AlphaFold2 model, which limited precision and yielded nine non-tolerant sites (Fig. 2d). The final 3 sites occurred in mTq2 (Fig. 2h), where the structurally constrained β-barrel fold is expected to exhibit position-dependent insertion sensitivity.

The remaining 10 intolerant sites mapped to Gpa1 but exhibited a distinct phenotype, reflecting a defining feature of this system. In the yeast mating pathway, signaling is driven primarily by the Gβγ subunit, whereas in many mammalian GPCR systems, Gα stimulates downstream signaling by producing second messengers. In this context, Gpa1 sequesters Gβγ to suppress basal activity, such that disruption of this interaction by peptide insertion can lead to signaling in the absence of an α-factor agonist.

Consistent with this model, insertion chimeras at 10 of 19 Gpa1 sites exhibited substantial basal signaling above baseline (Fig. 2f), indicating altered sequestration of Gα-Gβγ. These effects spanned a continuum of basal activity, demonstrating that insertion position differentially modulates the extent of disruption. Despite elevated basal signaling, some chimeras remained responsive to agonist (Fig. 2e), indicating that pathway function is retained despite partial loss of regulation at the Gα-Gβγ interface.

Together, these results show that peptide insertions are broadly tolerated across the yeast mating pathway, with 63% of sites retaining measurable signaling (Fig. 2i). Tolerance varies by protein: Ste2, Fus3, and mTq2 accommodate insertions at most positions, whereas intolerance clusters in Ste4, Ste12, and Gpa1. Within tolerated sites, insertion position modulates pathway output across a continuum rather than simply preserving or abolishing function. In Gpa1, many insertions disrupt Gα–Gβγ sequestration and induce basal signaling, while remaining responsive to agonist stimulation. These results demonstrate that it is possible to explore chimera space at the pathway scale to identify insertion sites that install new functionality while preserving core protein function.

### Global mapping of in-cell protection across the yeast mating pathway

The concept of protection reflects how structure and macromolecular assembly govern the accessibility of specific protein sites. Given the stable, yet dynamic nature of protein structure, buried or sterically occluded sites are less frequently exposed, whereas flexible or solvent-exposed sites are more accessible. Classical approaches such as hydrogen–deuterium exchange^33–35^, pulse proteolysis^36^, and fast quantitative cysteine reactivity^37,38^ can be used to probe and quantify these properties, but are limited to purified proteins.

Having shown that peptide insertions were broadly tolerated across the mating pathway, we expanded our chimera library to address this challenge and measure in-cell protection. Each HCV peptide insertion strain was paired with a companion strain expressing both a chimera and the cognate HCV NS3/4A protease, yielding a total of 182 CRISPR-engineered strains. This matched insertion-protease design enables each engineered site to report local accessibility within its native context and, to our knowledge, establishes the first pathway-scale measurement of site-specific protection in living cells.

As illustrated in Fig. 3a, proteolytic cleavage can produce two opposing responses. In one case, cleavage reduces or abolishes pathway activity (Fig. 3a, left). In the other, cleavage relieves an inhibitory constraint, specifically Gα-mediated sequestration of Gβγ, and causes or elevates basal signaling (Fig. 3a, right). Ste2 exemplifies the first behavior, where cleavage of the insertion site disables receptor function and reduces agonist-stimulated output. Gpa1 exemplifies the second behavior, where cleavage disrupts the Gα–Gβγ interaction, thereby releasing Ste4 and elevating basal signaling, an effect suppressed by protease inhibition.

**Fig. 3.**
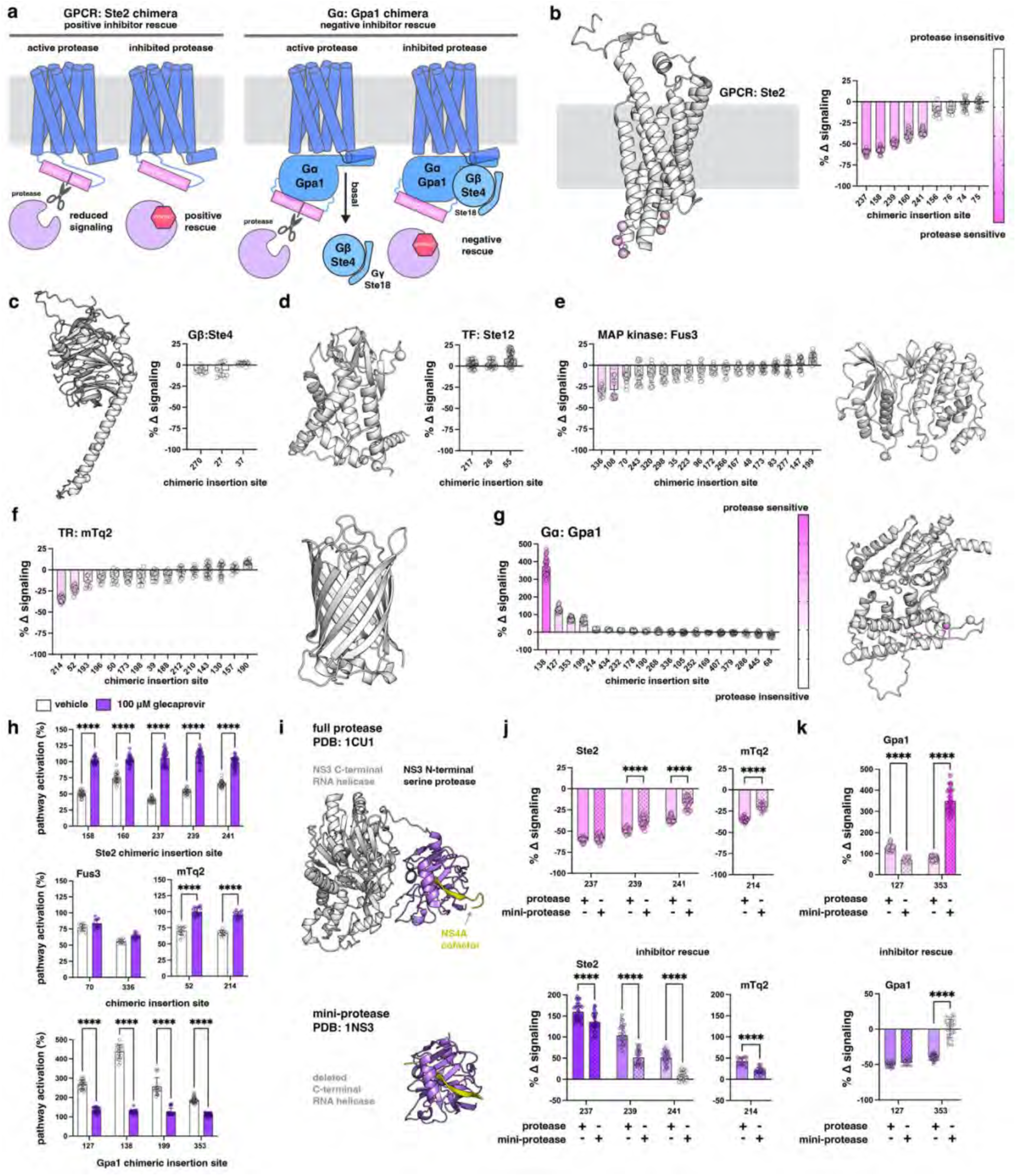
Global mapping of in-cell protection across the yeast mating pathway. **a**, Schematic of protease-dependent modulation of pathway signaling. Cleavage by the HCV NS3/4A serine protease yields two outcomes: loss of α-factor–dependent signaling with restoration upon inhibition (positive rescue, left), or induction of signaling upon cleavage with loss upon inhibition (negative rescue, right). **b-g**, Protease-dependent changes in signaling (± protease) for functional chimeras measured in the presence (**b-f**) or absence (**g**) of α-factor. Spheres (structural models) and bars (graphs) are color-coded from dark pink (protease-sensitive) to white (insensitive). **h**, Inhibitor rescue of protease-sensitive Ste2, Fus3, mTq2, and Gpa1 chimeras treated with glecaprevir (purple) or vehicle (white). **i**, Structural models of full-length (PDB: 1CU1) and truncated mini-protease (PDB: 1NS3) of HCV NS3/4A. The mini-protease comprises the NS3 serine protease domain (purple) lacking the RNA helicase domain (grey); the NS4A cofactor is shown in yellow. **j**,**k**, Comparison of full-length and mini-protease activity. Top, percent change in signaling in the presence (**j**) or absence (**k**) of 10 μM α-factor. Bottom, inhibitor rescue with 100 μM glecaprevir in the presence (**j**) or absence (**k**) of 10 μM α-factor. Cultures were normalized and incubated for 18–24 h. Data represent mean ± s.d. of ≥3 independent biological replicates, each with four technical replicates. Statistical analysis was performed using two-way ANOVA (**P ≤ 0.01, ***P ≤ 0.001, ****P ≤ 0.0001). See Supplementary Fig. S1–16 for additional data.

As shown in Fig. 3b–g, most of the 57 tolerant chimeras (Fig. 2i) were protected from protease cleavage, indicating their peptide inserts are occluded within their native structural or complex contexts. Protection was widespread across components, including Fus3 (16 of 18 sites; Fig. 3e) and mTq2 (13 of 15 sites; Fig. 3f), as well as all tolerated sites in Ste4 and Ste12 (Fig. 3c,d). In contrast, Ste2 showed more limited protection, with 4 of 9 sites protected, consistent with greater accessibility of its intracellular loops (Fig. 3b). Similarly, in Gpa1, 4 of 9 sites that did not exhibit substantial basal signaling (Fig. 2e) exhibited limited protection (Fig. 3g), as protease cleavage at these positions caused basal signaling.

Having established that proteolysis alters signaling at less protected sites, we next tried to reverse these effects by treating cells with the clinically approved NS3/4A inhibitor glecaprevir^39,40^ (Fig. 3h). In this framework, inhibitor treatment should suppress or prevent cleavage to restore chimera function, yielding either positive (Fig. 3a, left) or negative (Fig. 3a, right) rescue of pathway output. As shown in Fig. 3h, glecaprevir restored signaling at all protease-sensitive sites in Ste2 and mTq2 (positive rescue) and robustly suppressed protease-induced basal signaling by restoring Gpa1 sequestration of Gβγ (negative rescue).

In contrast, Fus3 sites could not be rescued (Fig. 3h), indicating a distinct mechanism of protection. Constitutive cleavage of Fus3 is consistent with an EX1-like regime, in which sites are persistently exposed and efficiently cleaved^33–37^, limiting recovery of protein function. In contrast, Ste2, mTq2, and Gpa1 exhibited inhibitor-responsive behavior consistent with an EX2-like regime^33–37^, in which sites dynamically alternate between protected and exposed states and are cleaved intermittently. In this regime, inhibitor treatment restores signaling by shifting the population toward intact protein.

In addition to structural accessibility, probe size is a key determinant of protection, with smaller probes more effectively accessing occluded sites. To test this in vivo, we repeated proteolysis and rescue experiments for select chimeras using a truncated version of the HCV NS3/4A protease. The N-terminal domain of NS3 encodes the serine protease activity, stabilized by the NS4A cofactor, whereas the C-terminal domain contains a DExH/D-box RNA helicase (Fig. 3i). Removal of this helicase domain generates a smaller mini-protease that retains catalytic activity^41^.

Using this mini-protease, we assessed site protection as a function of enzyme size in Ste2 and Gpa1. As shown in Figs. 3j–k, the mini-protease produced modest but consistent additional reductions in signaling at Ste2 insertion sites (239 and 241) and at an mTq2 site (214) compared with the full-length protease (Fig. 3h), indicating enhanced access to these recognition sites. Inhibitor treatment restored signaling when the full-length protease was expressed, but rescue was diminished with the mini-protease and was not observed at Ste2 site 241. These results are consistent with increased access shifting these sites from an EX2-like regime with the full-length protease to an EX1-like regime with the mini-protease.

Performing this analysis on select Gpa1 sites (Fig. 3j), we observed two distinct outcomes. At site 353, the mini-protease produced greater basal signaling than the full-length protease, and inhibitor treatment suppressed basal signaling only with the full-length enzyme, consistent with enhanced cleavage by the smaller probe. This behavior further indicates that increased accessibility shifts this site from an EX2-like regime with the full-length protease to an EX1-like regime with the mini-protease. In contrast, site 127 showed comparable inhibitor rescue with both protease forms, indicating that cleavage at this position is less sensitive to enzyme size.

Together, these results show that in-cell biophysical protection can be resolved at site resolution across a signaling pathway by integrating proteolysis, inhibition, and probe size. These measurements distinguish EX1- and EX2-like accessibility regimes in living cells, linking structural context to functional outcomes. By leveraging PAM scanning to generate a pathway-wide library of insertion sites, this approach enables systematic interrogation of protection in native protein and complex environments. These findings extend principles established in vitro to living cells and provide a generalizable strategy for incorporating peptide-recognition sequences into proteins for tool development, custom enzyme engineering, and monitoring viral protease evolution and escape.

### Engineering bi-functional yeast GPCR Ste2 chimeras

For our next demonstration of PAM scanning as a generalizable engineering strategy, we engineered bi-functional GPCR chimeras that incorporate reporter proteins while retaining receptor signaling. As shown in Fig. 4a, using the yeast GPCR Ste2 as a model, we inserted fluorescent (mNeonGreen) and luminescent (NanoLuc) reporters at 10 positions distributed across the N-terminus and extracellular loop 1 (ECL1). Of the ten insertion sites tested, three produced bi-functional chimeras that mapped to positions within ECL1, indicating a localized region of structural permissiveness. Focusing on these bi-functional chimeras, we found that signaling output was reduced relative to wild-type Ste2, whereas reporter protein function was robust and comparable across all permissive insertion sites.

**Fig. 4.**
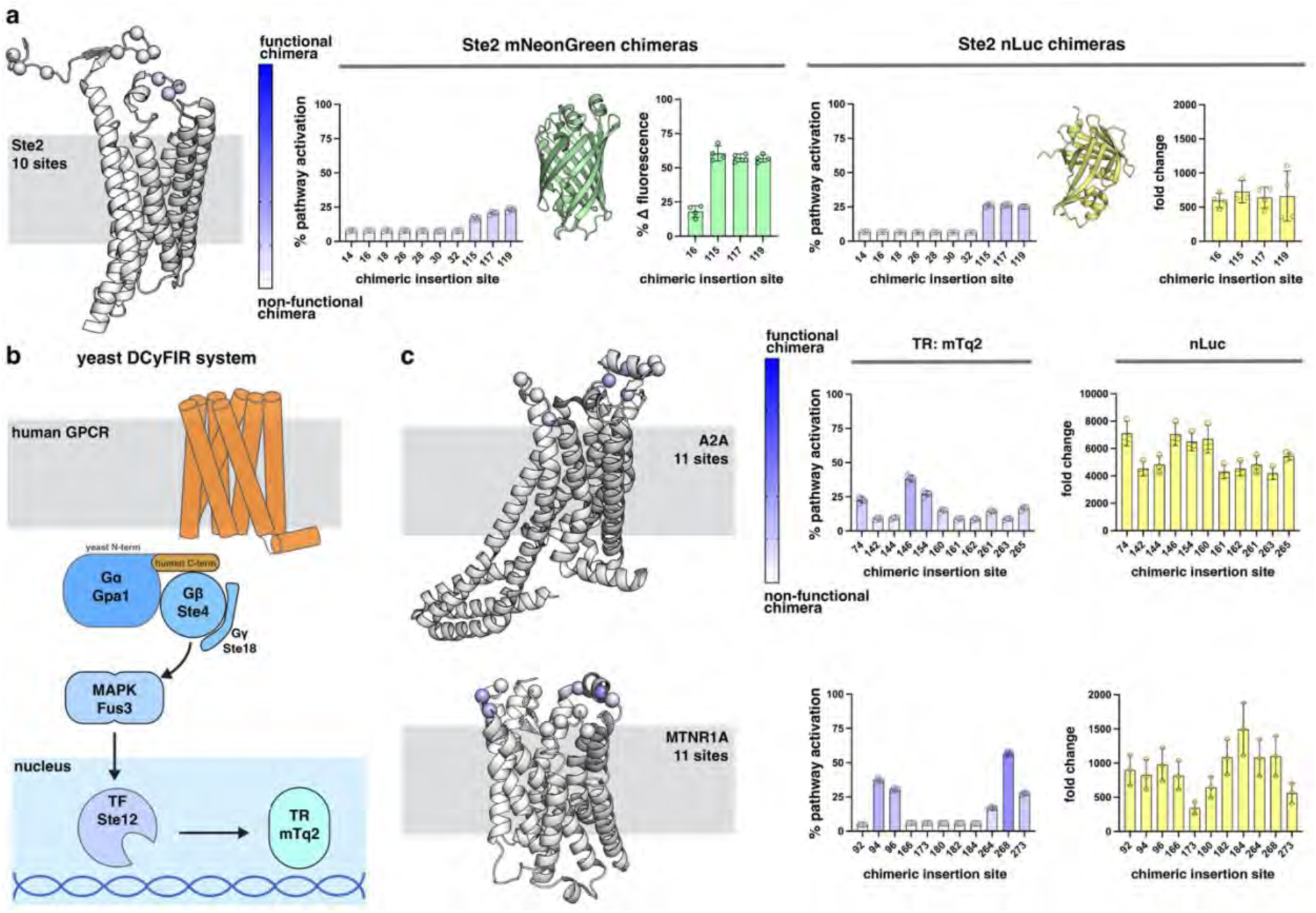
Engineering bi-functional GPCR chimeras. **a**, Functional screen of yeast GPCR chimeras containing inserted mNeonGreen (left) or NanoLuc luciferase (nLuc; right) reporters. **b**, Schematic of the DCyFIR platform adapted for human GPCR signaling in yeast. A chimeric Gα subunit couples human GPCR activation to the yeast pathway by replacing the five C-terminal residues of yeast Gα with those from human Gα (here, Gαi). Pathway activation is measured by endpoint fluorescence of the mTq2 transcriptional reporter. **c**, Functional screen of human A2A and MTNR1A receptor chimeras containing nLuc inserts. Signaling was stimulated with 1 mM adenosine (A2A) or melatonin (MTNR1A). Reporter output (mTq2 fluorescence or nLuc luminescence) and pathway activation were measured in parallel from separate aliquots after 18–24 h incubation in the absence of ligand unless otherwise indicated. Data are reported relative to WT as mean ± s.d. (n = 3 biological replicates). See Methods for details.

For non-permissive insertion sites, the absence of signaling did not necessarily reflect loss of Ste2 folding or reporter function. To confirm this distinction, we further investigated the insertion site 16 chimera as a representative example. As shown in Fig. 4a, although this chimera failed to signal, it retained detectable, albeit reduced, mNeonGreen fluorescence and NanoLuc luminescence levels comparable to bi-functional chimeras. These observations indicate that the site 16 chimera is folded and can support reporter protein function despite its signaling defect. Given that inserting one protein sequence into another is expected to substantially perturb local structure, this outcome is likely common across candidate insertion sites, underscoring the necessity of PAM scanning to systematically identify positions that support true bi-functional chimera engineering.

### Engineering bi-functional human GPCR chimeras

Having demonstrated proof of concept using Ste2, we next turned to the translational engineering of bi-functional human GPCRs. To this end, we employed our previously established DCyFIR platform^26–31^, a yeast-based system for functional interrogation of human GPCR signaling. As illustrated in Fig. 4b, human GPCRs are expressed in yeast and coupled to a re-engineered mating pathway in which receptor activation engages a chimeric Gα protein (Gpa1) composed of a human C-terminus fused to a yeast N-terminus, enabling productive coupling of human GPCRs to the endogenous yeast signaling machinery. Signal propagation proceeds through the yeast Gβγ subunit Ste4 and the MAP kinase Fus3, culminating in activation of the transcription factor Ste12 and induction of a fluorescent reporter (mTq2). This system enables quantification of human GPCR signaling and provides a robust framework for extending PAM scanning to bi-functional human GPCR engineering.

Using the DCyFIR platform, we applied PAM scanning to two representative human GPCRs, the adenosine A2A receptor (A2A) and the melatonin receptor MTNR1A. As part of the DCyFIR workflow, we CRISPR-integrated each receptor into the genome prior to scanning. We then performed PAM scanning by inserting NanoLuc at 11 extracellular positions (Fig. 4c, left) and quantified mTq2 signal output alongside NanoLuc luminescence (Fig. 4c, right). Notably, all A2A chimeras retained measurable signaling, albeit with varying magnitudes, indicating broad tolerance for extracellular insertions in this receptor. In contrast, only five MTNR1A chimeras preserved detectable signaling, highlighting more stringent structural constraints. Across both receptors, reporter activity was robust at most insertion sites, even when signaling was compromised, indicating that loss of pathway activation more often reflects disruption of receptor coupling rather than failure of chimera folding or stability.

Building on these yeast-based screening results, the DCyFIR platform provides a practical route for translational GPCR engineering by enabling rapid, scalable prioritization of candidate bi-functional chimeras. This approach allows insertion sites that preserve receptor signaling and reporter function to be efficiently advanced into more labor-intensive validation studies in human cell systems. To translate these findings into a human cellular context, we evaluated the subset of bi-functional chimeras identified by DCyFIR using a cAMP-based GloSensor assay in HEK293T cells (Fig. 5a). In this assay, intracellular cAMP binding promotes assembly of a split firefly luciferase, generating a luminescent signal that quantitatively reports GPCR-dependent regulation of cAMP levels. Accordingly, activation of Gs-coupled receptors increases cAMP production and elevates GloSensor luminescence, whereas activation of Gi-coupled receptors inhibits cAMP formation and results in decreased luminescence.

**Fig. 5.**
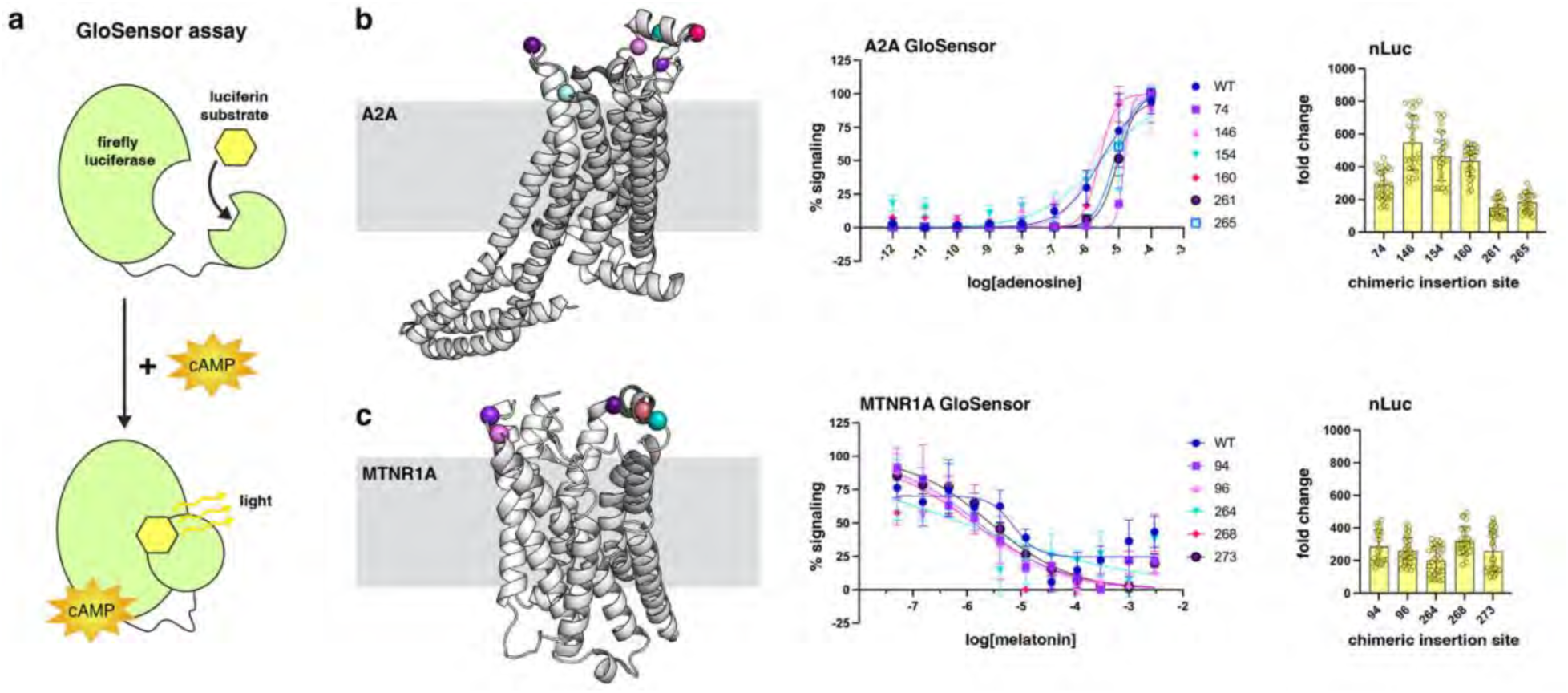
Validating yeast-screened bi-functional GPCR-reporter chimeras in human cells. **a**, Schematic of the GloSensor cAMP assay used to quantify GPCR-stimulated cyclic AMP (cAMP) signaling. **b,c**, Functional validation of A2A adenosine receptor–nLuc (**b**) and MTNR1A melatonin receptor–nLuc (**c**) chimeras in HEK293T cells. Left, structural models with nLuc insertion sites indicated. Middle, dose–response curves assessing GPCR signaling. Right, endpoint nLuc luminescence confirming reporter function. Insertion sites (left) are color-coded to match the corresponding datasets (middle and right). Data represent mean ± s.d. of three biological replicates, each with ≥3 technical replicates.

Focusing on the subset of A2A and MTNR1A chimeras identified as bi-functional by DCyFIR, we used the GloSensor assay to confirm bi-functional activity in human cells. As shown in Fig. 5b for the Gs-coupled A2A receptor, each bi-functional chimera exhibited a positive, adenosine-dependent dose–response curve comparable to wild-type A2A, indicating preservation of ligand-dependent activation and coupling to cAMP production. In contrast, as shown in Fig. 5c for the Gi-coupled MTNR1A receptor, each bi-functional chimera exhibited a negative, melatonin-dependent dose–response curve, again similar to wild-type MTNR1A, consistent with ligand-dependent inhibition of cAMP formation. Across both receptor sets, reporter activity remained robust and readily detectable, confirming maintenance of bi-functional behavior following translation from yeast to human cells.

## Discussion

PAM scanning enables systematic exploration of chimera-permissive sites at a scale difficult to achieve with conventional protein engineering approaches. Because any gene expressible in yeast can be subjected to CRISPR-directed insertion, this framework is broadly applicable to proteins from across all domains of life. Engineering at the DNA level enables rapid, parallel generation of insertion variants with minimal cost and labor compared to iterative protein-level design. Although we used SpCas9 here, the approach is readily extendable to alternative CRISPR systems with distinct PAM requirements, further expanding the accessible sequence space for insertional scanning. In this way, PAM availability is a programmable feature that defines a tunable landscape for systematic chimera design.

The ability to map permissive insertion sites in living cells enables a wide range of applications. Insertion-based reporters and biosensors can be rapidly developed by placing functional domains at sites that preserve host protein activity while reporting conformational or signaling changes. This approach is particularly well-suited for pharmacologically important targets such as GPCRs, where insertion into intracellular or extracellular loops enables real-time measurement of receptor activation or localization and can serve as fiducial markers to facilitate cryoEM studies. More broadly, systematic identification of permissive sites may accelerate the development of engineered proteins for research and therapeutic applications by reducing the need for trial-and-error design.

By combining insertional scanning with site-specific proteolysis, we identified chimeras that remain functional yet are responsive to protease cleavage, enabling reversible control of protein activity in living cells. Protease inhibition restores signaling at these sites, establishing a tunable system in which function can be modulated by cleavage and rescue. This creates two immediate applications. First, using optimized insertion sites, protease activity and inhibition can be monitored in pooled experiments, enabling rapid assessment of the effectiveness of existing inhibitors against naturally occurring or engineered viral protease variants. Second, this system can be used to screen libraries of peptide recognition motifs against a given protease or, conversely, to evolve proteases with desired specificity against defined peptide sequences.

Lastly, advances in sequence- and structure-based AI models have outpaced the availability of scalable experimental systems for testing their predictions. PAM scanning provides a direct route to close this gap by enabling computational design of insertion libraries and rapid experimental validation in living cells. While current models can estimate structural features, they remain limited in predicting insertion tolerance and functional outcomes. By enabling systematic, site-resolved measurements of insertion tolerance and accessibility, PAM scanning establishes a source of ground truth for benchmarking and refining predictive models, enabling iterative cycles of prediction and validation. In this way, PAM scanning couples design and experiment, establishing a scalable framework for data-driven discovery and predictive chimera engineering.

## Author contributions

D.G.I. performed the informatics analyses, developed all of the code, managed the study, and wrote the manuscript. J.A., K.L., N.K., S.T., and B.C. performed all experiments.

## Competing interests

None

### Acknowledgements

This work was supported by the National Institutes of Health through an R35 Maximizing Investigators’ Research Award (R35GM119518 to D.G.I.), which provided core funding for this study. Additional support was provided by a Pap Corps Champions for Cancer Research Endowed Chair to D.G.I. and the Sylvester Comprehensive Cancer Center.

## PAM-scanning chimera design and validation

### Computational pipeline and software implementation

Insertional scanning was performed using a custom Python-based workflow, the PAM-Scanning Chimera Designer, developed to identify PAM-accessible positions and design synonymous silencing edits across open reading frames (ORFs). The software comprises three integrated modules: (1) a Tkinter-based graphical interface for input specification, (2) a pipeline controller that orchestrates file processing, PAM detection, and BLAST filtering, and (3) a core library implementing codon-level scanning, synonymous mutagenesis, and primer generation.

The GUI accepts five required input files: (i) the ORF sequence (ATG to stop), (ii) an extended ORF+ FASTA containing ±1000 bp genomic context, (iii) the host genome FASTA (used as the local BLAST+ database), (iv) a codon usage table (in .cusp or .txt format), and (v) an optional codon-selection sheet listing specific residues for targeted scanning. The user also provides the gene name, BLAST database label, and primer suffixes, along with numerical parameters defining primer length, the maximum allowed PAM–cut gap, the codon-sampling stride, and the off-target thresholds.

Upon execution, the program reads the input FASTA sequences and constructs a codon index across the ORF. Using a codon-usage table derived from the host organism, the algorithm maps each amino acid to its synonymous codons. The extended ORF+ sequence is then scanned on both strands for Cas9 recognition motifs (NGG and CCN), identifying PAM sites, associated guide sequences, and predicted Cas9 cut positions.

### Synonymous with PAM and guide silencing

To prevent re-cutting of successfully integrated constructs, each PAM site is subjected to synonymous “silencing” edits that remove Cas9 recognition without altering the encoded amino acid. Codons overlapping or flanking the PAM triplet are identified, and synonymous substitutions are introduced to eliminate the GG or CC dinucleotide motif. When the PAM lies out of frame or spans two codons, both codons are evaluated for permissible substitutions based on codon-usage bias.

If direct PAM silencing is not possible, the pipeline performs guide-sequence silencing by introducing synonymous substitutions within the 20-nucleotide protospacer to increase mismatches to the original guide while preserving the protein sequence. This fallback ensures that each site yields a viable “safeguide,” a modified target resistant to Cas9 cutting after chimeric integration.

### Off-target filtering and guide selection

All candidate guides are screened for potential off-target alignments using local NCBI BLAST+ against the specified host genome. Alignments are parsed to identify “PAM inclusions,” defined as partial matches exceeding user-defined identity or alignment length thresholds. Guides with significant off-target similarity are excluded.

Remaining safeguide candidates are ranked according to two criteria: (1) the distance between the predicted Cas9 cut site and the targeted codon (PAM–cut gap) and (2) the number of PAM inclusions in the host genome. The optimal guide per codon is selected based on the minimal gap and the lowest off-target burden.

### Primer generation and library design

For each optimal safeguide, oligonucleotide primers flanking the insertion site are designed with user-defined lengths and suffix sequences compatible with the cloning backbone used for Cas9-mediated integration. Primers are automatically organized into 96- or 384-well plate layouts to facilitate large-scale oligonucleotide ordering.

To assess target coverage, the pipeline computes the fraction of the ORF reachable by valid PAM sites within the maximum allowed cut distance. Output directories include:

- **QC/** — raw and silenced FASTA files,
- **BLAST+/** — off-target analysis reports,
- **ORDER/** — primer order spreadsheets, and
- **WARNINGS/** — flagged codons with residual PAM inclusions.

### Chimera construction in yeast

Chimeric constructs were generated by CRISPR-mediated homologous recombination in *Saccharomyces cerevisiae*. For each PAM-scannable position, the designed primer pair was used to amplify and insert a cassette encoding a protease cleavage site or reporter in-frame at the safeguide locus. Integration events were selected on defined media and verified by colony PCR and Sanger sequencing across both junctions. Details of CRISPR transformation, payload preparation, and functional validation are provided below.

## Summary of experimental reagents

All primers, plasmids, strains, and reagents used or created in this study are listed in Datasets S1-4.

## Plasmid construction

### CRISPR PAM-scan Plasmid Construction

CRISPR plasmids were constructed using the multicopy pML104 X-2 UnTS vector previously developed in our laboratory^1–4^. This plasmid contains a 2-micron origin of replication, encodes Cas9 and a single-guide RNA (sgRNA) scaffold with a 20-base-pair unique targeting sequence (UnTS), and confers URA3 selection in yeast. For each chimera, the UnTS was replaced with a 20-base-pair safeguide sequence targeting a specific codon insertion site. These safeguide sequences were generated using our custom computational pipeline and software implementation described above. The safeguide was introduced into the pML104 X-2 UnTS backbone using a chimera-specific forward primer (annotated as gF in Dataset S1) and a universal reverse primer carrying a 5′ phosphate modification (pML guide UR (5PHOS): 5′-phos-GATCATT TATCTT TCACTG CGGA-3′).

Plasmid assembly and circularization were performed using the Around-the-World PCR protocol. Each reaction contained 1.25 µL of guide (gF) primer (5 µM), 1.25 µL of pML guide UR (5PHOS) primer (5 µM), 9.0 µL of nuclease-free water, 12.5 µL of Q5 2× Master Mix (New England Biolabs, M0492L), and 1.0 µL of template plasmid DNA (10 ng/µL). The presence of the PCR product was confirmed by agarose gel electrophoresis before adding 1.0 µL of DpnI (NEB, R0176S) to digest methylated template DNA. The mixture was incubated at 37 °C for 1.5 hours, then heated to 80 °C for 20 minutes, and finally held at 4 °C. To ligate the PCR-amplified plasmid, 23.0 µL of nuclease-free water, 5 µL of 10× T4 DNA ligase buffer (NEB, B0202S), and 1.0 µL of T4 DNA ligase (NEB, M0202L) were added, and the reaction was incubated overnight at 16 °C. Ligation was terminated by heating at 65 °C for 10 minutes, and the newly assembled plasmids were subsequently transformed into bacteria as described below.

### HiFi Plasmid Construction

For plasmid assembly using HiFi cloning, we PCR-amplified both the plasmid backbone and corresponding insert using HiFi-compatible primers designed for sequence overlap. Following PCR, the template DNA was digested with DpnI (NEB, R0176S) to remove methylated parental plasmid. The HiFi assembly reaction was prepared by combining 14.2 µL of nuclease-free water, 0.5 µL of insert PCR product, 0.3 µL of vector PCR product, and 5 µL of HiFi DNA Assembly Master Mix (New England Biolabs, E2621L) in a single PCR tube, followed by incubation at 50 °C for 15 minutes. The assembled plasmids were then introduced into *E. coli* (DH5α) by transformation using 2 µL of the final HiFi reaction for amplification and sequence verification, as described in the protocol below.

### Protease Plasmid Construction

The gene encoding the Hepatitis C virus NS3/NS4A protease was obtained from Addgene (plasmid #61696) in a pETDuet bacterial expression backbone. Using HiFi DNA assembly, we subcloned the protease coding sequence into the pYE181 yeast vector under the control of the TEF1 promoter and CYC1 terminator using primers pYE MCS F and R (Dataset S1). The resulting plasmid was not used for direct expression of the protease in yeast but served as a storage and amplification vector to provide template DNA for subsequent CRISPR-mediated integration of the TEF1-protease-CYC1 open reading frame into the yeast genome.

### Mini Protease Plasmid Construction

A truncated version of the Hepatitis C virus NS3/NS4A protease, containing only the N-terminal NS3 serine protease domain, and missing the C-terminal RNA helicase domain, was generated using HiFi DNA assembly. The mini protease coding sequence^5^ was subcloned into the pYE181 yeast vector under the control of the TEF1 promoter and CYC1 terminator using the primers pYE181.hifi.fwd.ns34a.construct and ns34a.construct.hifi.rvs (Dataset S1). As with the full-length protease plasmid described above, the resulting construct was not used for direct mini protease expression in yeast but served as a storage and amplification vector to provide template DNA for CRISPR-mediated integration of the TEF1–protease–CYC1 open reading frame into the yeast genome.

### Chimeric GPCR Plasmid Construction

Human GPCR chimera plasmids were generated by PCR amplification of the chimeric A2A and MTNR1A receptor genes that had been previously integrated into the yeast genome via CRISPR editing. Amplification was performed using primer pairs A2A pcDNA3.1 (pos) F/R and MTNR1A pcDNA3.1 (pos) F/R (Dataset S1). The resulting PCR products were then subcloned into the pcDNA3.1(+) expression vector using the HiFi plasmid construction procedure described above, enabling subsequent expression of the chimeric GPCRs in human cell lines.

## CRISPR payload design and preparation

### CRISPR payload design

Yeast CRISPR reactions were performed by transforming a plasmid expressing a single-guide RNA (sgRNA), the Cas9 endonuclease, and a URA3 selectable marker, as described above. Expression of these components generates a precise double-strand break at the genomic locus complementary to the sgRNA target sequence. The payload refers to the repair DNA fragment introduced during this process, consisting of the desired gene or nucleotide sequence flanked by homology arms corresponding to the genomic regions immediately upstream and downstream of the cut site. In PAM-scanning experiments, the payload encoded a peptide or protein sequence, which was inserted into a target gene to generate a chimeric construct. The sequences of the homology arms and the corresponding primers for payload amplification were determined computationally using the custom pipeline described above.

### CRISPR Payload Preparation for Integrating Protease Recognition Sites

Because the Hepatitis C virus NS3/4A protease recognition sequence (EDVVCCSMSY) is encoded by only 30 nucleotides (5′-GAAGATGTTGTCTGTTGCTCTATGTCATAT-3′), we designed the CRISPR payload as complementary 100-base-pair forward and reverse primers (Dataset S1). To generate double-stranded DNA, equal volumes of each primer (15 µL) were combined with 30 µL of Q5 2× Master Mix (New England Biolabs) in a PCR tube. The mixture was heated in a thermocycler at 95 °C for 1.5 minutes, followed by a gradual cooling step of 1 °C per minute until the temperature reached 50 °C to allow primer annealing.

### CRISPR Payload Preparation for Integrating Reporter Proteins

For reporter proteins nLuc and mNeonGreen, CRISPR payloads were generated using a two-step PCR-based process. In the first step, each reporter gene was amplified from its respective storage plasmid or gblock (pNluc-N1, from Lambert Lab; mNeonGreen, IDT-DNA) using primers that appended universal flexible linkers: an N-terminal SGGSGG linker (5′-TCAGGTGGCAGTGGAGGT-3′) and a C-terminal GGSGGS linker (5’- AGACCCGCCGCTACCTCC-3’). In the second step, the amplified reporter constructs were re-amplified using 100-base-pair primers containing homology arms specific to the designated chimeric insertion site, thereby generating the final payloads for yeast genome integration.

## General molecular biology protocols

### Bacterial Transformation

Chemically competent E. coli DH5α cells (a gift from the Dohlman laboratory) were prepared and stored at –80 °C using the Mix & Go! E. coli Transformation Kit (Zymo Research, T3002). For transformation, 50 µL of competent cells were thawed on ice for 20 minutes, mixed with 2 µL of synthesized plasmid DNA, and incubated on ice for an additional 5 minutes. The transformation mixture was then spread onto LB agar plates containing ampicillin (0.01%) and incubated overnight at 37 °C, with plates positioned inverted during incubation.

### Plasmid Isolation and Verification

Three individual bacterial colonies were each inoculated into 3 mL of fresh LB broth containing ampicillin (0.01%) in 15 mL Falcon conical tubes and incubated overnight at 37 °C in a shaking incubator. Plasmid DNA was isolated the following day using the EZ BioResearch Miniprep Kit (M1000-250) according to the manufacturer’s protocol. The purified plasmids were sequence-verified by Sanger sequencing (Eurofins Genomics) before downstream applications.

### Yeast Transformation

To prepare a starter culture, 3 mL of YPD medium was inoculated with yeast and incubated overnight at 30 °C with shaking. The following day, 100 µL of the overnight culture was transferred into 5 mL of fresh YPD and incubated under the same conditions for approximately 2.5 hours, until the culture reached an OD₆₀₀ of 0.6–1.2. Cells were harvested by centrifugation, and the supernatant was discarded. The yeast pellet was sequentially washed with 5 mL of 1× TE buffer and 5 mL of LiOAc mix, pelleting after each wash. Following the final wash, the pellet was resuspended in 200 µL of LiOAc mix, sufficient for approximately four transformations.

Each transformation reaction combined 50 µL of chemically prepared yeast cells with 300 ng of plasmid DNA (e.g., CRISPR guide plasmid), 20 µL of repair DNA or donor payload, 350 µL of PEG mix, and 5 µL of salmon sperm DNA (Invitrogen, 15632011) in a sterile 1.5 mL microcentrifuge tube (USA Scientific, 1615-5500). Samples were vortexed briefly and incubated at room temperature for 30 minutes. After incubation, 24 µL of 100% DMSO was added, and the cells were subjected to a heat shock at 42 °C for 15 minutes. After heat shock, cells were pelleted by centrifugation (10,000 rpm, 3 minutes), gently resuspended in 100 µL of YPD (without vortexing), and plated onto appropriate selective media (e.g., –Ura plates) for approximately 3 days at 30 °C. Unless otherwise noted, all yeast pelleting steps were performed by centrifugation at 3,000 × g for 3 minutes.

### Yeast Strain Verification

3-12 yeast colonies were inoculated into individual wells of a 96-deep-well block, each containing 1 mL of –Ura selection medium. The block was sealed with a breathable film (Thomas Scientific, 1198M9), briefly shaken at 1,400 rpm for 1 minute, and then incubated without shaking for approximately 2 days at 30 °C. Following incubation, chimeric integrations were verified using the genomic DNA (gDNA) extraction and verification protocol described below. Confirmed chimeras were streaked onto 5-fluoroorotic acid (5-FOA) plates and incubated for 2–3 days at 30 °C.

From each 5-FOA plate, three colonies were selected and reinoculated into fresh wells of a 96-deep-well block containing 1 mL of YPD medium per well. The block was incubated overnight at 30 °C, after which the gDNA verification protocol was repeated to confirm the strain identity. Verified YPD cultures were prepared for long-term storage by mixing 300 µL of culture with 130 µL of 50% glycerol (Sigma-Aldrich, G6279-1L) and storing the glycerol stocks at –80 °C.

## Yeast media

### Yeast Extract Peptone Dextrose Media (YPD)

Yeast extract peptone (YP) solution was prepared by combining 20 g/L peptone (RPI, P20240-1000.0), 10 g/L yeast extract (RPI, Y20020-1000.0), and 900 mL of double-distilled water (ddH₂O), then autoclaved. Immediately before use, 100 mL of sterile 20% (w/v) D-glucose (RPI, G32045-3000.0) was added to the YP solution to yield complete YPD medium.

To prepare YPD agar plates, the same formulation was used, except that D-glucose (20 g/L) and agar (15 g/L; RPI, A20030-1000.0) were added directly to the mixture before autoclaving. The sterilized medium was then poured into 100 mm sterile Petri dishes (VWR, 25384-342) and allowed to solidify before use.

### Synthetic Complete Dropout (SCD) Medium (Dropout Media)

Uracil dropout medium (–U) was prepared by combining 5.00 g/L ammonium sulfate (Sigma-Aldrich, A4418-500G), 1.70 g/L yeast nitrogen base without amino acids or ammonium sulfate (RPI, Y20060-250), one NaOH bead pellet (ACS, BDH9292-500G), 20 g/L D-glucose (2% w/v), and 0.77 g/L CSM–Ura supplement (MP Biomedicals, 4511222). The medium was sterilized by vacuum filtration (VWR, 10040-440).

For uracil selection plates, the same formulation was used, except that agar (15 g/L) was added before autoclaving to sterilize the mixture. The molten medium was then poured into 100 mm sterile Petri dishes and allowed to solidify before use.

### Low Fluorescence SCD Media (LoFo)

Low-fluorescence medium was prepared by combining 8.72 g/L potassium phosphate dibasic (Sigma-Aldrich, P3786-2.5kg), 9.76 g/L MES hydrate (RPI, M22040-500.0), 5.00 g/L ammonium sulfate (Sigma-Aldrich, A4418-500G), 1.70 g/L yeast nitrogen base—low fluorescence (without amino acids, folic acid, or riboflavin; Formedium, CYN6510), 0.79 g/L CSM Dropout Mix (complete) (MP Biomedicals, 4500022), and 20.0 g/L D-glucose (2% w/v). The solution pH was adjusted to the desired value before filter sterilization. The prepared medium was stored in the dark and allowed to stabilize for approximately 3 days before use in fluorescence-based assays.

### 5-FoA selection plates

A 100 mM 5-FOA stock solution was prepared by dissolving 5 g of 5-fluoroorotic acid (5-FOA) (Zymo Research, F9001-5) in 50 mL of DMSO (Sigma-Aldrich, 472301-1L). The solution was incubated in a 42 °C water bath for approximately 15 minutes, or until the powder fully dissolved, then aliquoted into 10 mL portions and stored at –20 °C. To prepare 5-FOA selection plates, the following components were combined in 900 mL of double-distilled water (ddH₂O): 15 g agar, 20 g D-glucose, 5.00 g/L ammonium sulfate, 1.70 g yeast nitrogen base (without amino acids or ammonium sulfate), 0.79 g CSM complete mix, and 30 mg uracil (Sigma-Aldrich, U1128-100G). ddH₂O was added to a final volume of 1 L, and the mixture was autoclaved for sterilization. After cooling slightly, the 5-FOA stock solution was added to the molten medium, which was then poured into 100 mm sterile Petri dishes and allowed to solidify. It is important to note that 5-FOA plates are light sensitive and must be stored in the dark to preserve activity.

## Bacterial transformation media and agar plates

### Luria–Bertani (LB) Broth

LB broth was prepared by combining 10 g/L Bacto tryptone (RPI, T60060-500.0), 5 g/L Bacto yeast extract (RPI, Y20020-1000.0), 5 g/L sodium chloride (NaCl) (Sigma-Aldrich, 567440-1KG), and one NaOH pellet in distilled water. The mixture was stirred until all components were fully dissolved, adjusted to volume, and autoclaved to sterilize before use.

### LB-Ampicillin Broth and Plates

LB–Amp broth was prepared fresh for each experiment. The required volume of sterile LB broth was aliquoted into 15 mL (VWR, 21008-918) or 50 mL (VWR, 21008-951) conical tubes. A frozen stock solution of 100 mg/mL carbenicillin in 50% ethanol (AG Scientific, C-1385-25GM) was added to achieve a final antibiotic concentration of 0.1% (v/v)—for example, 3 µL of antibiotic stock per 3 mL of LB broth.

LB–Amp agar plates were prepared following the same protocol used for LB broth, except that 18 g/L Bacto agar was added before autoclaving. After sterilization, the molten medium was allowed to cool until warm to the touch, and carbenicillin was added to a final concentration of 100 µg/mL. The medium was then poured into 100 mm sterile Petri dishes and allowed to solidify at room temperature.

## Yeast transformation solutions

### 1× TE Buffer

A 10× TE buffer stock was prepared by dissolving 12.12 g/L Tris base (Sigma-Aldrich, 93352-1kg) and 2.92 g/L EDTA (Sigma-Aldrich, EDS-100G) in distilled water. The stock was diluted 1:10 with water to yield a 1× working solution containing 10 mM Tris and 1 mM EDTA.

### LiOAc Mix

A 10× (1 M) lithium acetate (LiOAc) stock was prepared by dissolving 102 g/L LiOAc (Sigma-Aldrich, 517992-100G) in water. To prepare the LiOAc Mix, 100 mL of 10× LiOAc stock, 100 mL of 10× TE buffer, and 800 mL of water were combined, yielding a final concentration of 10 mM Tris, 1 mM EDTA, and 100 mM LiOAc.

### PEG Mix

The PEG mixture was prepared by combining 40 g of PEG 3350 (Sigma Life Sciences, 95904-1kg-F), 10 mL of 1 M LiOAc, and 10 mL of 10× TE buffer. The final solution contained 10 mM Tris, 1 mM EDTA, 100 mM LiOAc, and 40% (w/v) PEG 3350.

## Engineering yeast base strains

The Saccharomyces cerevisiae strain BY4741 (MATa his3Δ1 leu2Δ0 met15Δ0 ura3Δ0) was used as the parental background for strain construction. This strain was modified to include two safe-locus expression cassettes, the X-2 landing pad (LP) and X-II5 landing pad (LP), previously developed in our laboratory^1–4^, resulting in the P22Δ strain. Using CRISPR/Cas9-mediated genome editing, we generated a companion protease strain in which the Hepatitis C virus NS3 protease was integrated into the X-2 LP (X-2: NS3) and its NS4A cofactor into the X-II5 LP (X-II5: NS4), producing the P22Δ–NS3–4A base strain. Complete genotypes of all strains used in this study are provided in Dataset S3.

## Engineering of yeast strains containing chimeras of signaling proteins

Yeast strains harboring chimeric signaling proteins were generated using the CRISPR-based PAM-scan system. All guide plasmids and CRISPR payloads were designed using the computational pipeline described above. Successful genomic integrations were verified by PCR amplification of genomic DNA with chimera-specific primers, and in selected cases, the resulting chimeric loci were sequenced for downstream applications. No CRISPR editing errors were detected in any of the sequenced constructs.

### Mating pathway chimera screen

Each mating pathway chimera was integrated into two companion yeast strains. To generate each chimera, the Hepatitis C virus NS3/4A protease recognition sequence (EDVVCCSMSY) was inserted in frame at selected codon positions within key mating pathway components in the genome. Integration was performed in both the P22Δ strain, which lacks the protease, and the P22Δ-NS3-4A strain, which expresses the HCV NS3 protease from the X-2 locus (X-2: NS3) together with its NS4A cofactor from the X-II5 locus (X-II5: NS4). The functional activity of each resulting chimera was subsequently evaluated using the assay protocol described below.

### Yeast Ste2 and Human GPCR reporter chimeras

Each yeast-human GPCR chimera was engineered using a two-step CRISPR integration process. In Step 1, using the P22Δ base strain, the desired GPCR genes were integrated into the X-2 landing pad, using the approach established in our previous DCyFIR studies^1–4^. In Step 2, the resulting strains from Step 1 served as the base strains for chimeric integrations, in which the nLuc and mNeonGreen reporter genes were CRISPR-integrated in frame at selected codon positions within each GPCR. The functional activity of all resulting chimeras was subsequently evaluated using the assay protocol described below.

## Functional validation of PAM-scan chimeras

### General culture conditions

Yeast P22Δ chimera strains were inoculated into individual wells of a 96-deep-well block containing 1 mL of low-fluorescence (LoFo) medium (pH 6.0) and incubated overnight at 30 °C without shaking. The following day, 40 µL of saturated culture was transferred to a black 384-well microplate (Greiner, 781096), shaken at 2,200 rpm for 1 minute, and the optical density (OD_₆₀₀_) was measured using a Clariostar plate reader. Saturated cultures were diluted to a final OD_₆₀₀_ of 0.3 in 500–750 µL of LoFo medium (pH 6.0) and incubated at 30 °C for approximately 5 hours, typically reaching an OD_₆₀₀_ of 0.6–1.2.

### Functional validation of yeast mating pathway chimeras

Approximately 1 hour before yeast cultures reached the desired optical density, 384-well assay plates were prepared. A stock solution of α-factor ligand (Genescript, RP01002) was thawed from frozen and prepared by dissolving the powdered peptide in nuclease-free water (nfH_2_O; RPI, 7732-18-5). Ligand concentration was determined using a NanoDrop spectrophotometer with the Protein A₂₈₀ protocol (“Other protein E & MW”) employing an extinction coefficient of 12,490 M⁻¹cm⁻¹ and a molecular weight of 1,683.96 Da. Working stocks of 100 µM and 10 µM α-factor were prepared in nfH_2_O. For each yeast strain, 4 µL of either 100 µM or 10 µM α-factor was dispensed into four replicate wells (n = 4), while vehicle control wells received 4 µL nfH_2_O.

After approximately 5 hours of culture growth, the yeast was measured for OD_₆₀₀_ and diluted in LoFo medium (pH 6.0) to a final OD_₆₀₀_ of 0.1. 36 µL of the diluted culture was seeded into each ligand-containing well (n=4 technical replicates), resulting in final α-factor concentrations of 10 µM or 1 µM. Plates were sealed with breathable film, shaken at 2,200 rpm for 1 minute, and incubated at 30 °C for 18–24 hours.

Fluorescence of the mTq2 transcriptional reporter was measured using a Clariostar plate reader and the following settings:

- Measurement type: Fluorescence (endpoint)
- Flashes per well: 10
- Excitation: 430–10 nm
- Dichroic filter: LP 458
- Emission: 482–16 nm
- Gain: 1300
- Focal height: 6.5 mm
- Settling time: 0.5 s
- Reading direction: Bidirectional (left to right, top to bottom)

### Functional validation of human GPCR chimeras

The functionality of human GPCR chimeras was evaluated using the GαZ DCyFIR yeast strain developed in our prior work^1–4^. The yeast GPCR chimeras Ste2–nLuc and Ste2–mNeonGreen were tested in the same P22Δ strain background used for protein–peptide chimeras. Ligands were applied at the following final concentrations: 1 mM melatonin (Sigma-Aldrich, M5250-1G) for MTNR1A–nLuc, 1 mM adenosine (Sigma-Aldrich, A9251-25G) for A2A–nLuc, and 10 µM α-factor for Ste2–nLuc and Ste2–mNeonGreen chimeras. Yeast cultures were incubated with their respective ligands for 18–24 hours at 30 °C, after which fluorescence of the mTq2 transcriptional reporter was measured using a Clariostar plate reader with the same settings as above.

In parallel, using independent yeast cultures, we confirmed the functionality of the nLuc and mNeonGreen reporter inserts. For nLuc validation, the Nano-Glo Luciferase Assay System (Promega) was added directly to the yeast culture at a final concentration of 0.2×, and luminescence was measured immediately using the Clariostar (emission filter: 470–80 nm). To assess mNeonGreen activity, fluorescence was recorded on the same instrument (excitation: 430–10 nm; dichroic filter: LP 458; emission: 482–16 nm).

## Protease Cleavage/Protection and Inhibitor Rescue Assays

### Assessment of Protease Cleavage

Functionally validated Hepatitis C protease recognition site chimeras were further characterized in two stages. In the first stage, we assessed whether the NS3/4A protease cleaved the chimeric form of the mating pathway signaling component, as indicated by a loss or reduction in signaling activity. Yeast cultures were prepared according to the general culture conditions described above, treated with an α-factor agonist, incubated at 30 °C for 18–24 hours, and analyzed for signaling output using the mTq2 transcriptional reporter on a Clariostar plate reader.

### Inhibitor Rescue Assay

In stage two, we evaluated whether cleavage-induced signaling loss could be rescued by the Hepatitis C protease inhibitor Glecaprevir (MedChem Express, HY-17634). Experiments were conducted under the same general culture conditions as above, except that 32 µL of yeast culture (OD₆₀₀ = 0.1) was used per well. Each well received 4 µL of 100 µM α-factor and 4 µL of Glecaprevir working stock at final concentrations of 1 µM, 10 µM, or 100 µM. Vehicle control wells contained 4 µL of 10% DMSO (Sigma-Aldrich, 472301-1L). All conditions were performed in technical quadruplicate (n = 4).

### Basal Signaling Gpa1 Chimeras

For Gpa1 chimeras with inserted-induced basal signaling, α-factor was omitted. Wells received 4 µL of nfH₂O in place of ligand, followed by 32 µL of yeast culture (OD₆₀₀ = 0.1). Final glecaprevir concentrations were 1, 10, and 100 µM.

### Preparing stock Glecaprevir solutions

Glecaprevir was reconstituted in 100% DMSO to 10 mM, stored at –20 °C, and thawed immediately before preparing working stocks, each containing 10% DMSO.

## Functional Characterization of Human GPCR–nLuc Chimeras in HEK293T Cells

### Mammalian Cell Transfection

On the first day of experimentation, HEK293T cells were seeded at a density of 3 × 10⁶ cells per 10-cm dish in complete Dulbecco’s Modified Eagle Medium (DMEM; ThermoFisher, 11995081) and incubated at 37 °C in a humidified 5% CO₂ atmosphere for 24 hours to reach approximately 80% confluency. On the following day, each dish was transfected with a total of 4 µg plasmid DNA, consisting of 2 µg of the GPCR plasmid and 2 µg of the pGloSensor-22F plasmid (Promega, GU174434). Transfections were performed using the TransIT transfection reagent (Mirus Bio, MIR 5405) at a 3 µL reagent:1 µg DNA ratio, following the manufacturer’s protocol. Cells were then incubated under standard culture conditions (37 °C, 5% CO₂) for 24 hours post-transfection.

### Assay Setup in 384-well Plates

Transfected cells were harvested by adding 1.5 mL of trypsin (Gibco) to each 10-cm dish and briefly incubating to facilitate detachment. Trypsinized cells were neutralized with complete medium, collected, and centrifuged to pellet. The supernatant was removed, and the pellet was resuspended in 14 mL of fresh DMEM. Using a multichannel pipette, 40 µL of cell suspension was plated per well into white 384-well plates (Greiner, 781098) at a final density of approximately 20,000–40,000 cells per well, with n = 3 replicates per condition. Plates were incubated for 24 hours at 37 °C and 5% CO₂ to allow cell adherence and recovery.

### Luciferin Loading

On the fourth day, the culture medium was aspirated from each well using a vacuum aspirator. Each well then received 20 µL of 1× D-luciferin solution (GoldBio, LucNA) prepared in HBSS (Gibco) supplemented with 20 mM HEPES (pH 7.4). Plates were incubated at 37 °C, 5% CO₂ for 1 hour to allow luciferase substrate uptake and signal stabilization before ligand addition.

### Assessment of Melatonin–nLuc Receptor Chimeras

Following luciferin loading, 10 µL of isoproterenol (ISO; Sigma-Aldrich, I5627-5G) was added to achieve a final concentration of 250 nM in a total reaction volume of 40 µL. Plates were incubated at room temperature (RT) for 15 minutes. A serial ligand dilution assay (1:3) was then performed by adding 10 µL of melatonin at various concentrations (n = 3 technical replicates) to each well, followed by an additional 15-minute incubation at RT. Luminescence was measured using a Clariostar plate reader under GloSensor protocol settings (measurement type: luminescence; interval: 1 s; bidirectional vertical reading, left to right, top to bottom).

### Assessment of Adenosine–nLuc Receptor Chimeras

For A2A–nLuc chimeras, ligand activity was evaluated using a 1:10 serial dilution assay. After luciferin loading, 10 µL of adenosine was added to each well, yielding a final well volume of 30 µL. Plates were incubated for 15 minutes at RT before luminescence measurement using the same Clariostar GloSensor protocol as described above.

### nLuc Signaling Assay

To evaluate the activity of nLuc in the human GPCR–nLuc chimeras, furimazine substrate (Promega, Nano-Glo Luciferase Assay) was diluted 100× to 1× working concentration. After performing the GPCR ligand assays, baseline luminescence was measured after ∼1 hour to confirm minimal background signal. Then, 10 µL of 1× furimazine was added to each well, resulting in final substrate concentrations of 0.20× for MTNR1A–nLuc and 0.25× for A2A–nLuc chimeras. Plates were shaken in the dark at 300 rpm for 2 minutes, and luminescence was immediately measured on the Clariostar using the nLuc protocol (measurement type: luminescence; interval: 1 s; emission: 482–16 nm; gain: 3,000; focal height: 3 mm; bidirectional horizontal reading, left to right, top to bottom).

## Supplemental Figures and Legends

**Fig. S1.**
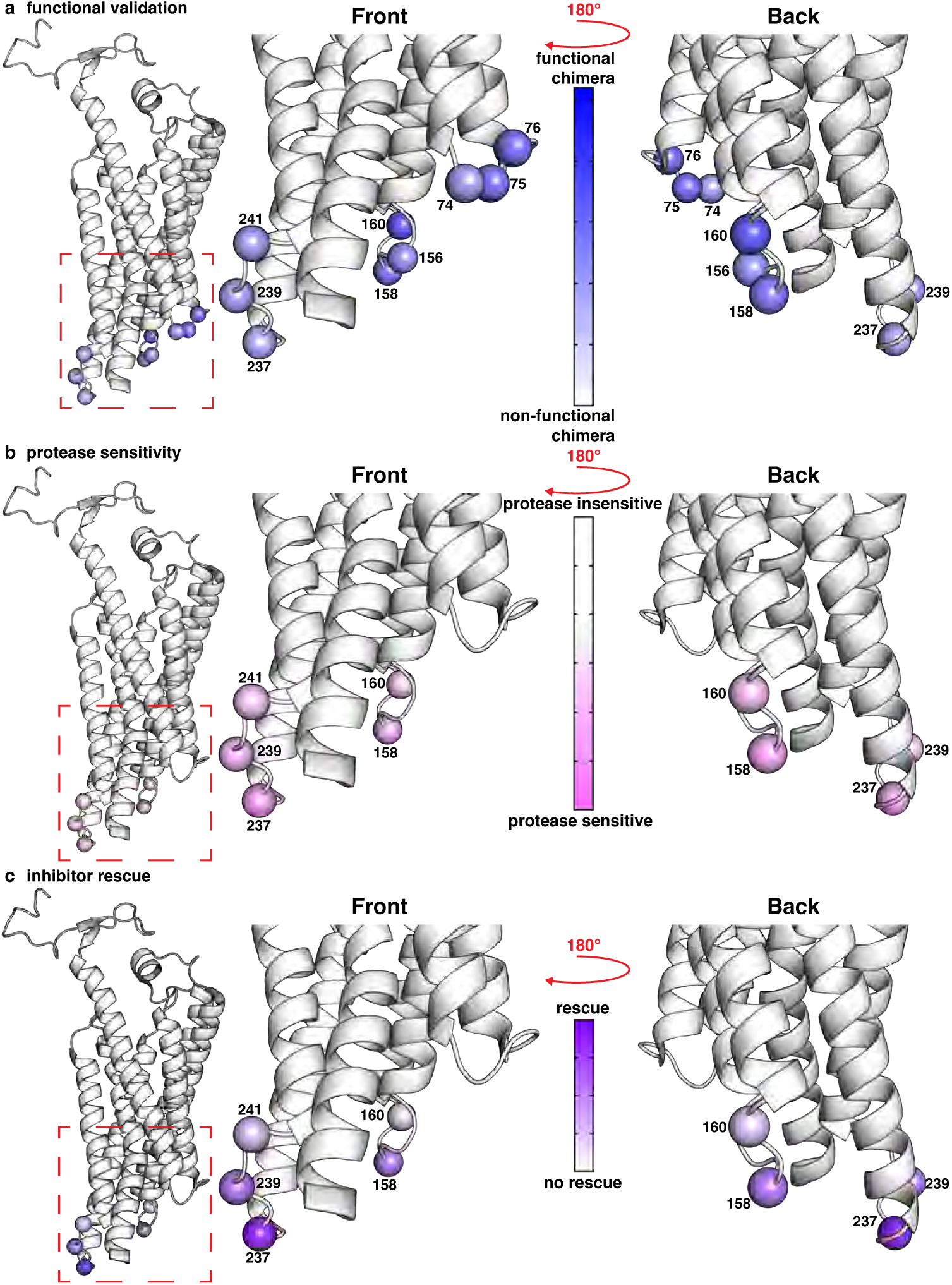
Ste2 HCV Peptide Insertion Sites. **a-c,** Structural views of Ste2 (PDB: 7AD3) with PAM-scan chimeric insertion positions labeled by codon number and represented as spheres.

**Fig. S2.**
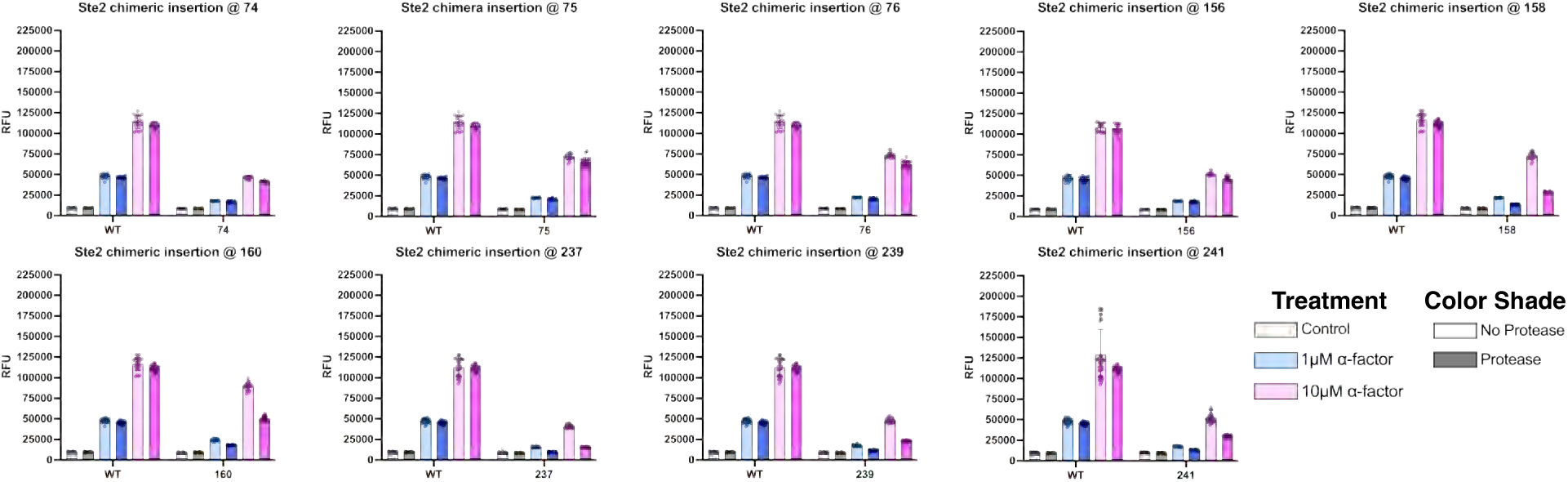
Primary functional validation data for Ste2 HCV peptide chimeras. Pheromone pathway activation was quantified by mTq2 fluorescence and reported as relative fluorescence units (RFU). Each condition includes a minimum of n = 3 independent experiments, each with n = 4 technical replicates.

**Fig. S3.**
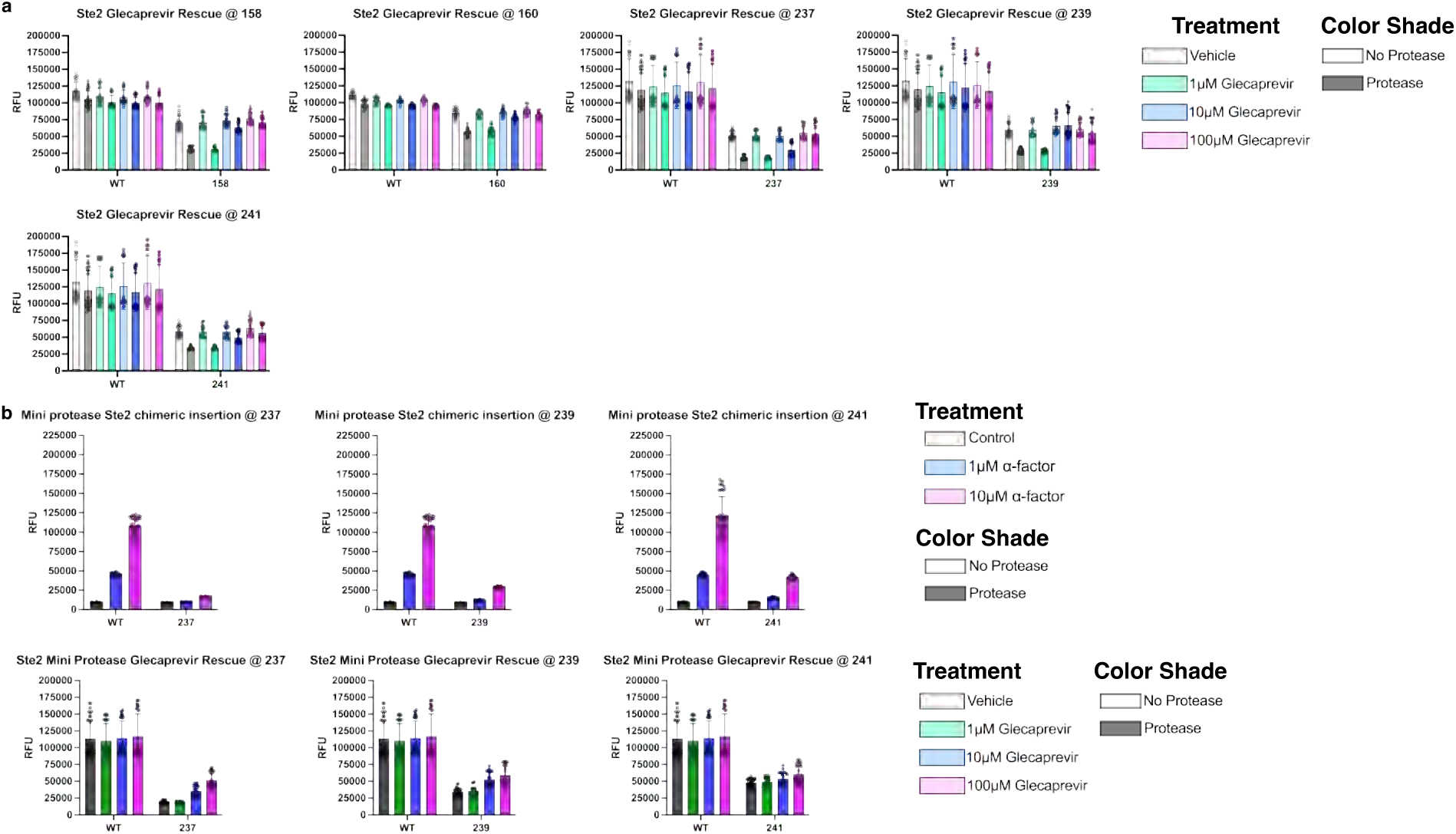
Primary functional validation, protection, and inhibitor rescue data for Ste2 HCV peptide chimeras. **a**, Primary data from the inhibitor assay performed on each Ste2 site that showed reduced protease-dependent signaling. **b**, Primary data from the mini-protease constructs. The α-factor assay (top) was used to derive protease activity, and the inhibitor assay (bottom) was used to calculate rescue by the inhibitor glecaprevir. Pheromone pathway activation was quantified by mTq2 fluorescence and reported as relative fluorescence units (RFU). Each condition includes a minimum of n = 3 independent experiments, each with n = 4 technical replicates.

**Fig. S4.**
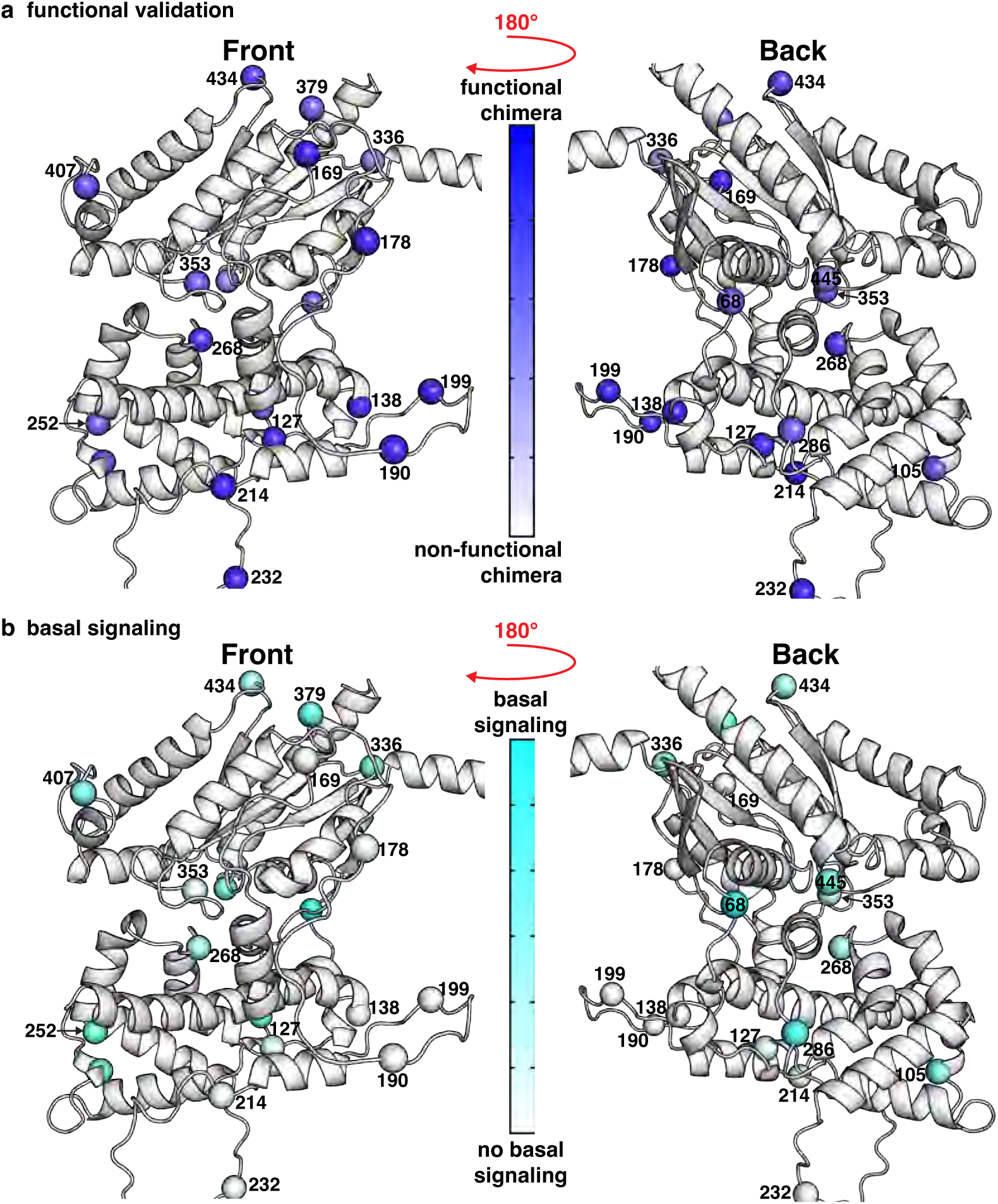
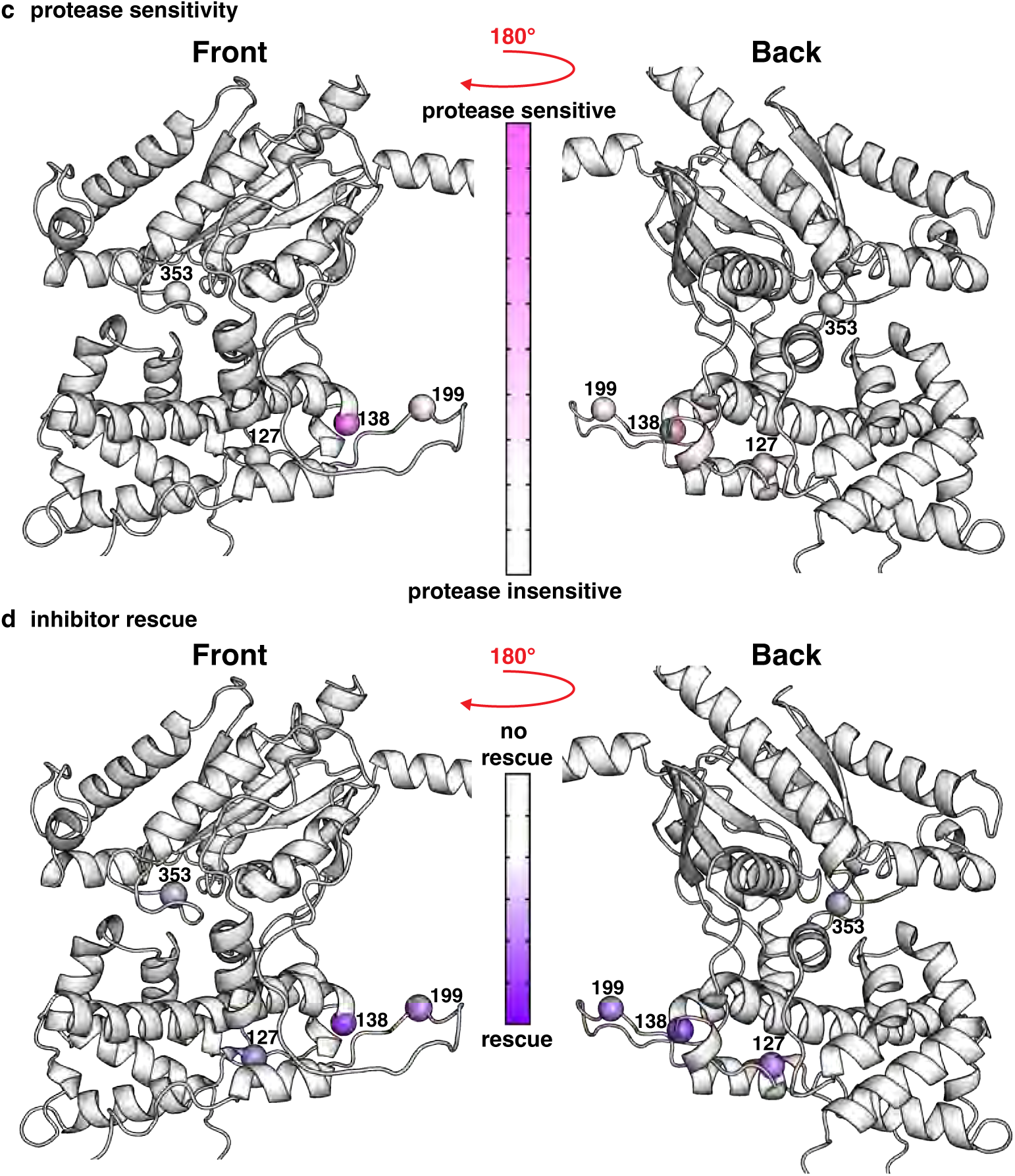
Gpa1 HCV Peptide Insertion Sites. **a-b,** Structural views of AlphaFold2 model of Gpa1 (AF-P08539-F1-model_v4-1) with PAM-scan chimeric insertion positions labeled by codon number and represented as spheres. **c-d,** Structural views of AlphaFold2 model of Gpa1 (AF-P08539-F1-model_v4-1) with PAM-scan chimeric insertion positions labeled by codon number and represented as spheres.

**Fig. S5.**
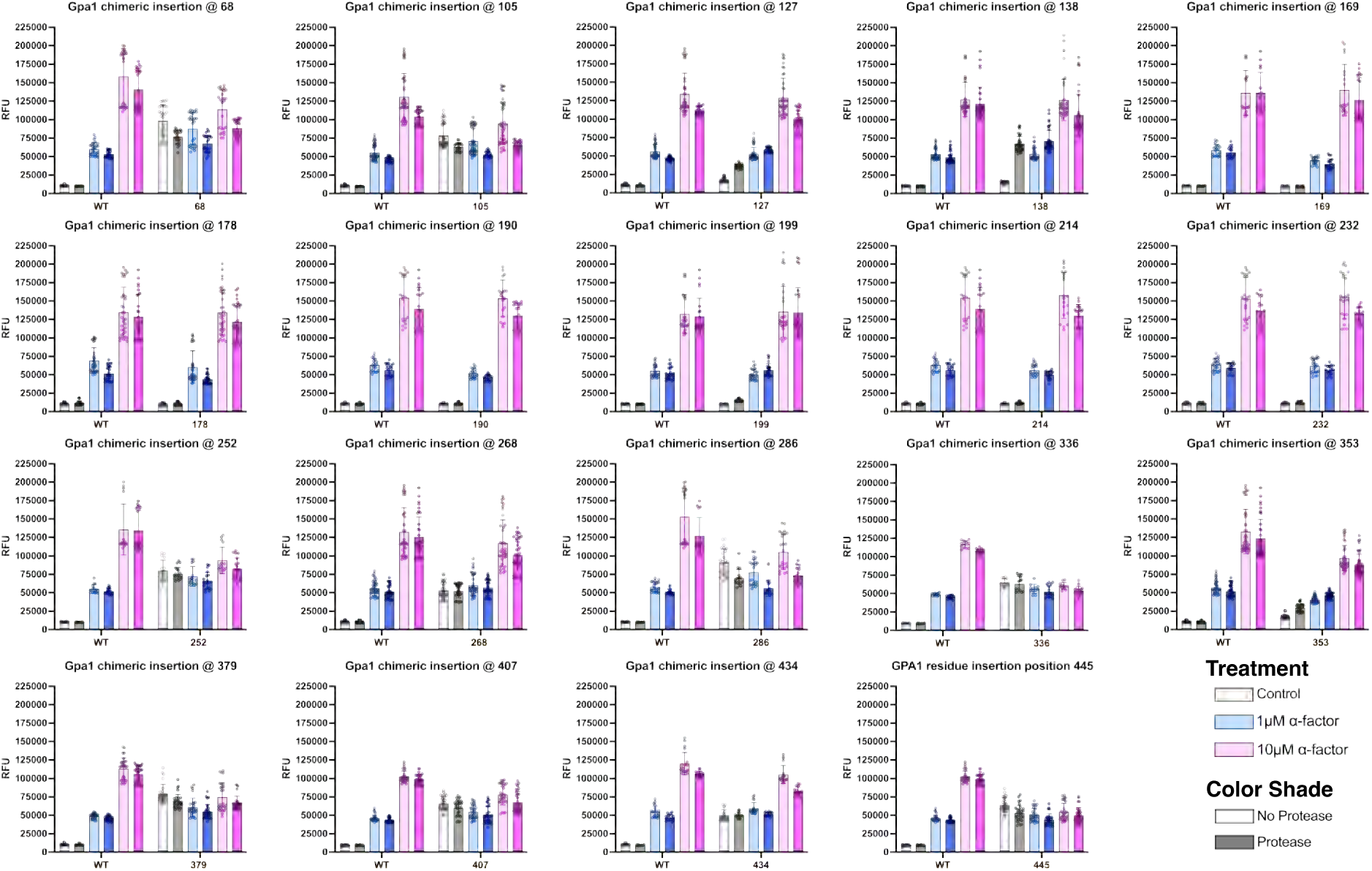
Primary functional validation data for Gpa1 HCV peptide chimeras. Pheromone pathway activation was quantified by mTq2 fluorescence and reported as relative fluorescence units (RFU). Each condition includes a minimum of n = 3 independent experiments, each with n = 4 technical replicates.

**Fig. S6.**
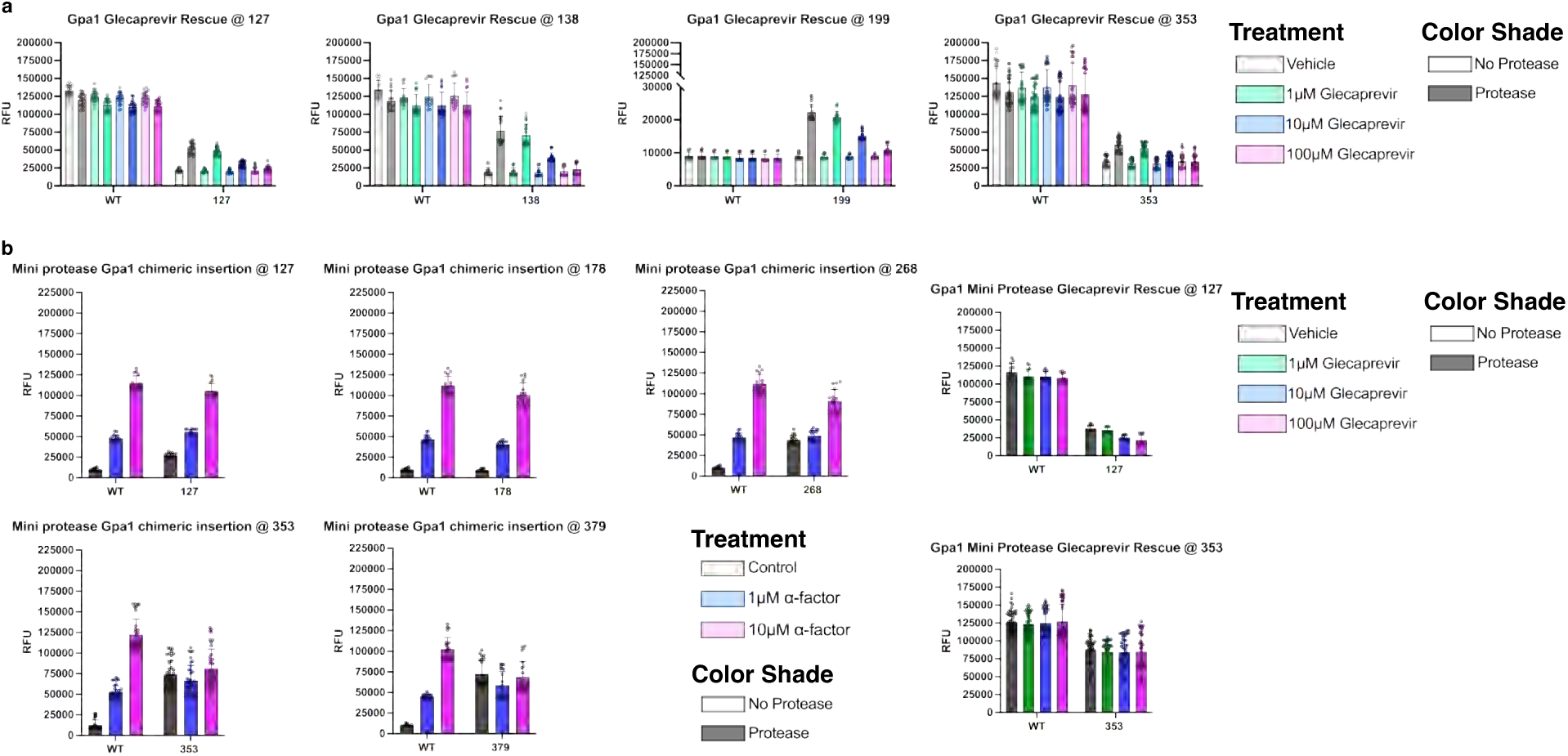
Primary functional validation, protection, and inhibitor rescue data for Gpa1 HCV peptide chimeras. **a**, Primary data from the inhibitor assay performed on each Gpa1 site that showed reduced protease-dependent signaling. **b**, Primary data from the mini-protease constructs. The α-factor assay (top) was used to derive protease activity, and the inhibitor assay (bottom) was used to calculate rescue by the inhibitor glecaprevir. Pheromone pathway activation was quantified by mTq2 fluorescence and reported as relative fluorescence units (RFU). Each condition includes a minimum of n = 3 independent experiments, each with n = 4 technical replicates.

**Fig. S7.**
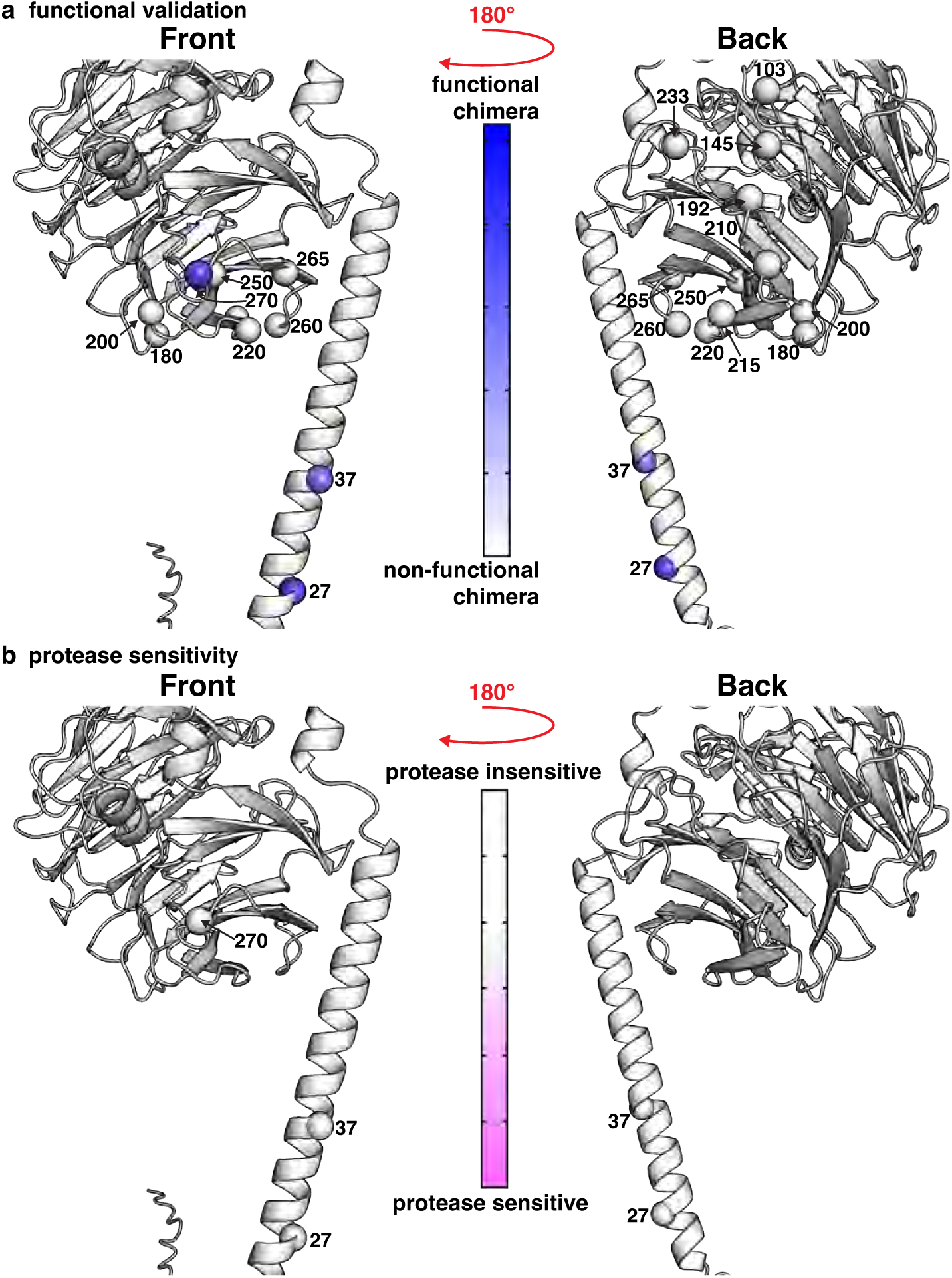
Ste4 HCV Peptide Insertion Sites. **a-b,** Structural views of AlphaFold2 model of Ste4 (AF-P18851-F1-model_v4) with PAM-scan chimeric insertion positions labeled by codon number and represented as spheres.

**Fig. S8.**
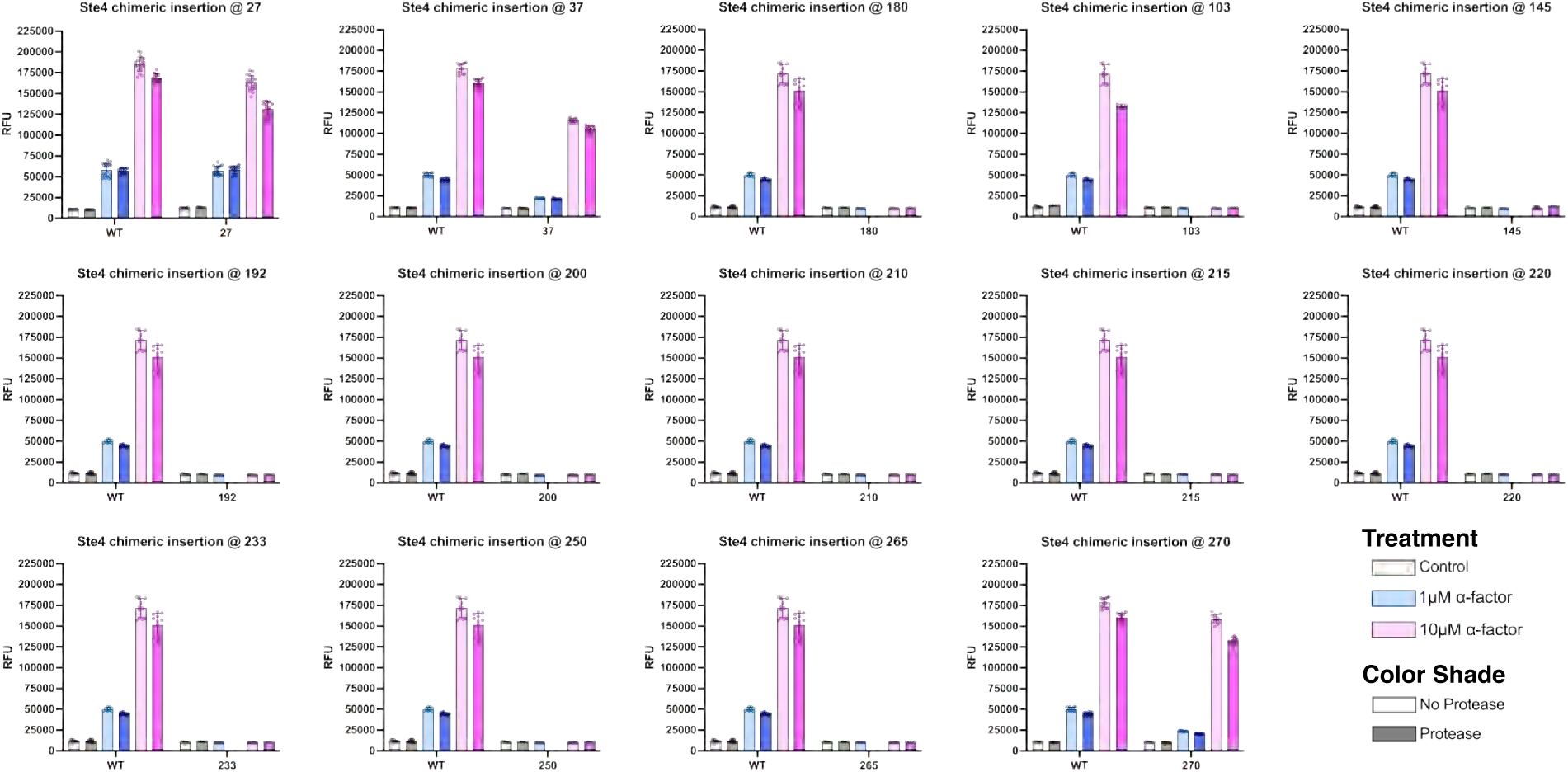
Primary functional validation data for Ste4 HCV peptide chimeras. Pheromone pathway activation was quantified by mTq2 fluorescence and reported as relative fluorescence units (RFU). Each condition includes a minimum of n = 3 independent experiments, each with n = 4 technical replicates.

**Fig. S9.**
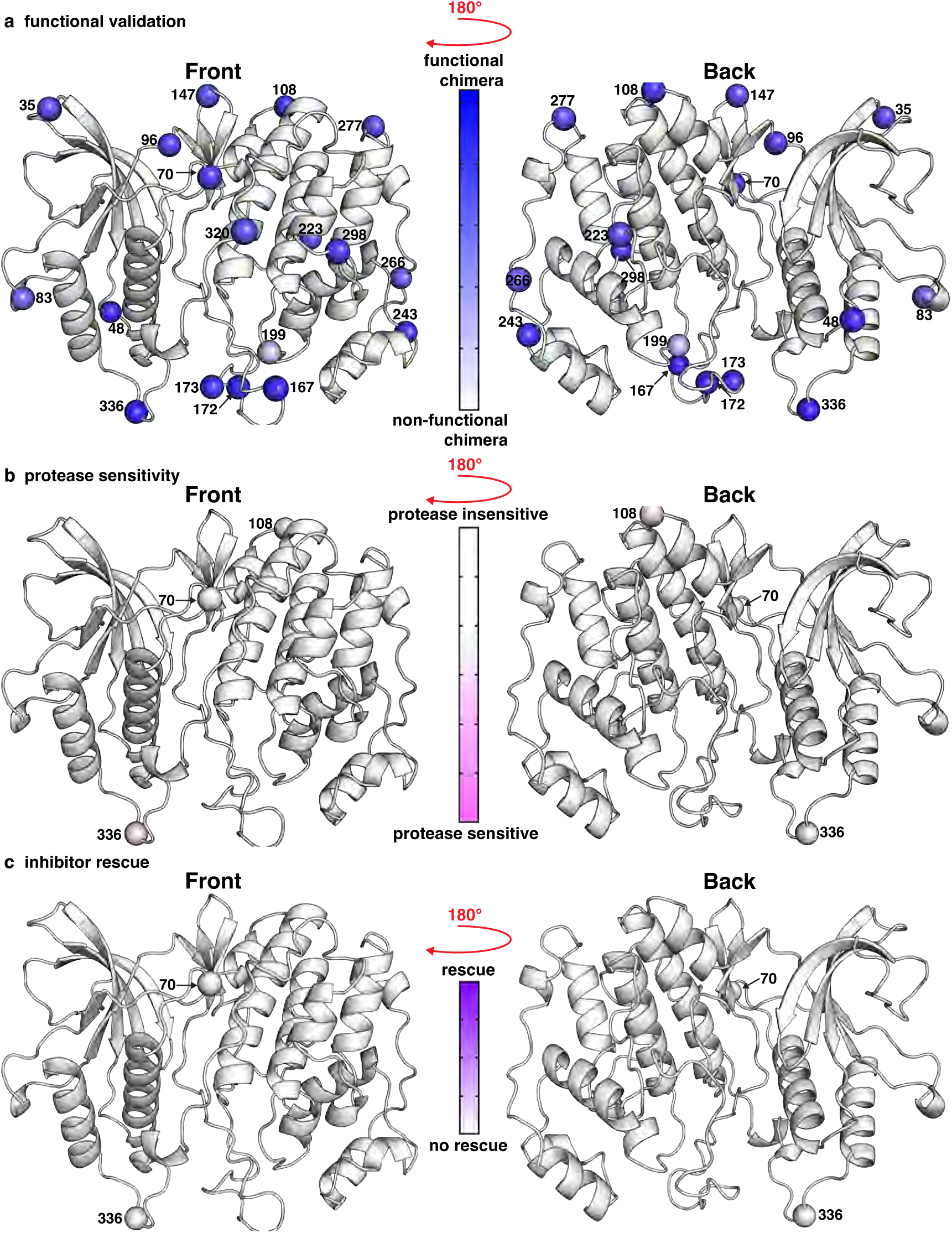
Fus3 HCV Peptide Insertion Sites. **a-b,** Structural views of AlphaFold2 model of Fus3 (AF-P16892-F1-model_v4) with PAM-scan chimeric insertion positions labeled by codon number and represented as spheres.

**Fig. S10.**
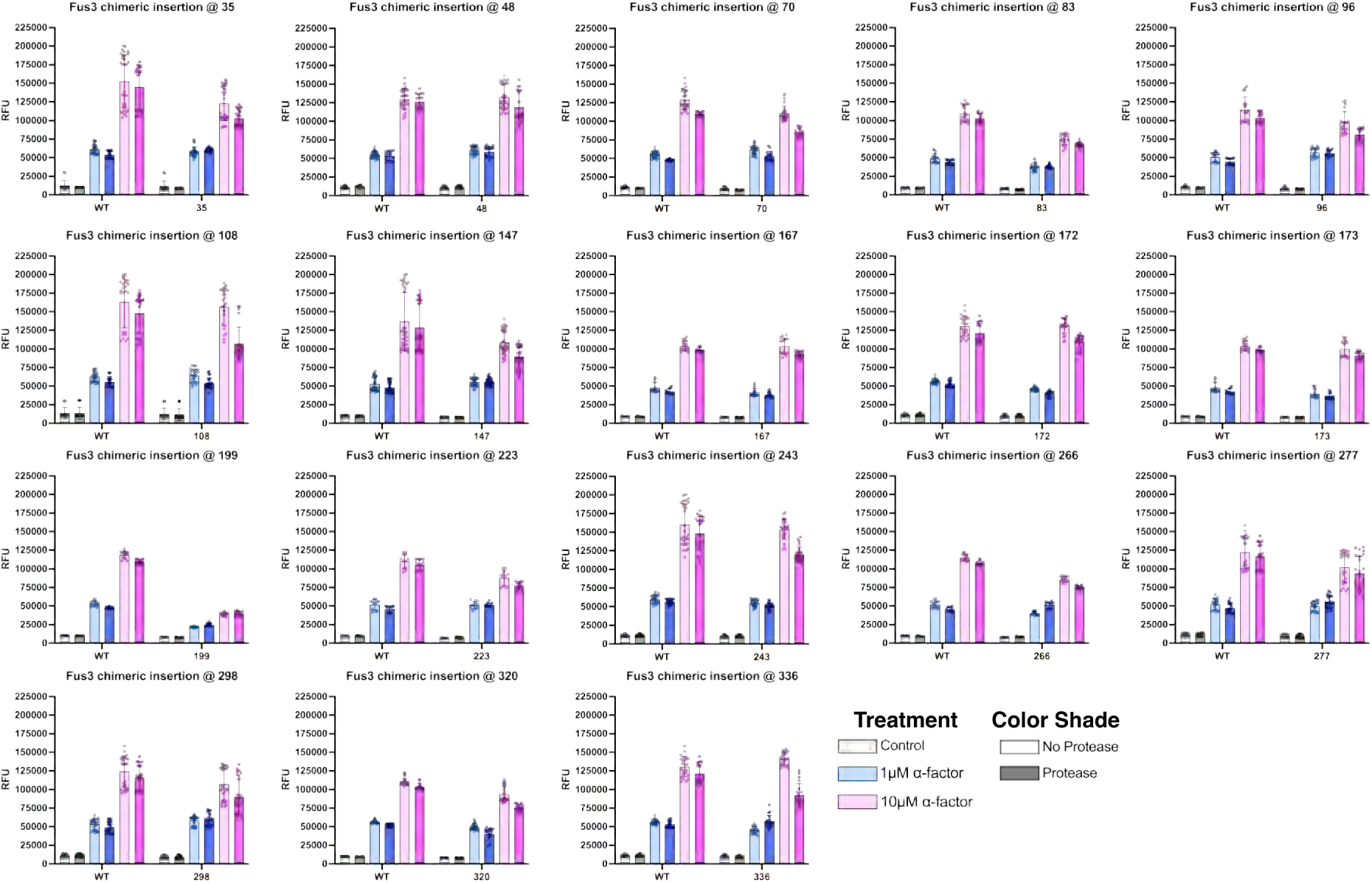
Primary functional validation data for Fus3 HCV peptide chimeras. **a**, Primary data from the inhibitor assay performed on each Fus3 site that showed reduced protease-dependent signaling. **b**, Primary data from the mini-protease constructs. The α-factor assay (top) was used to derive protease activity, and the inhibitor assay (bottom) was used to calculate rescue by the inhibitor glecaprevir. Pheromone pathway activation was quantified by mTq2 fluorescence and reported as relative fluorescence units (RFU). Each condition includes a minimum of n = 3 independent experiments, each with n = 4 technical replicates.

**Fig. S11.**
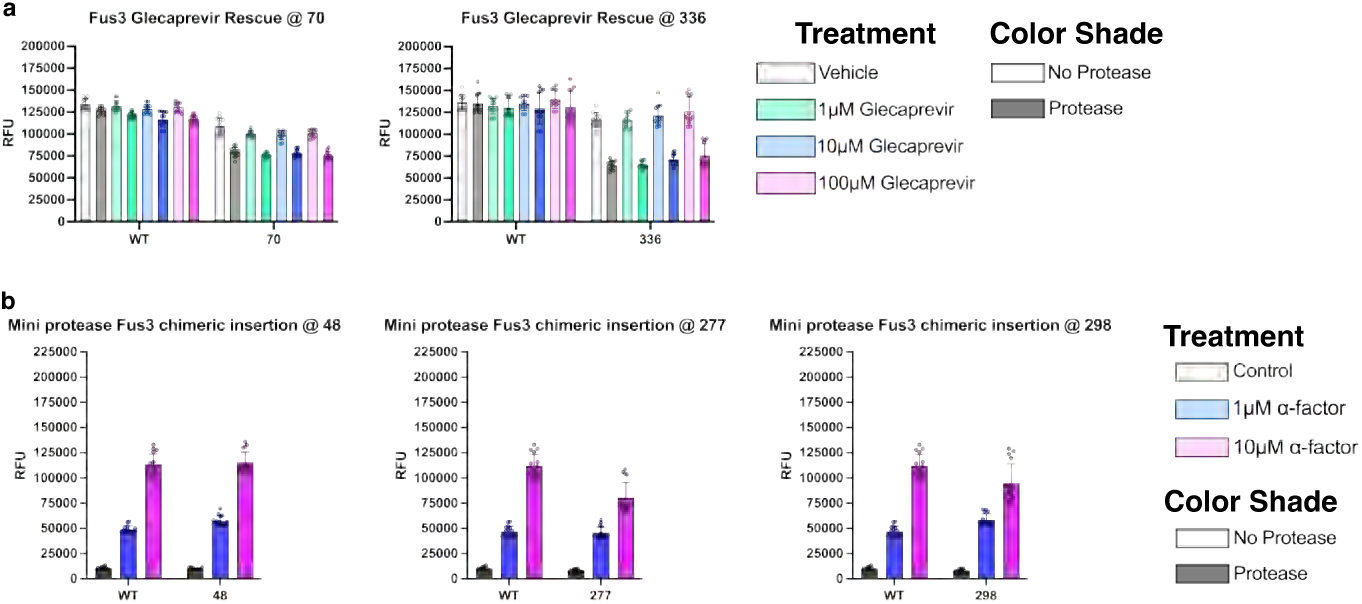
Primary functional validation, protection, and inhibitor rescue data for Fus3 HCV peptide chimeras. **a**, Primary data from the inhibitor assay performed on each Fus3 site that showed reduced protease-dependent signaling. **b**, Primary data from the mini-protease constructs. The α-factor assay (top) was used to derive protease activity, and the inhibitor assay (bottom) was used to calculate rescue by the inhibitor glecaprevir. Pheromone pathway activation was quantified by mTq2 fluorescence and reported as relative fluorescence units (RFU). Each condition includes a minimum of n = 3 independent experiments, each with n = 4 technical replicates.

**Fig. S12.**
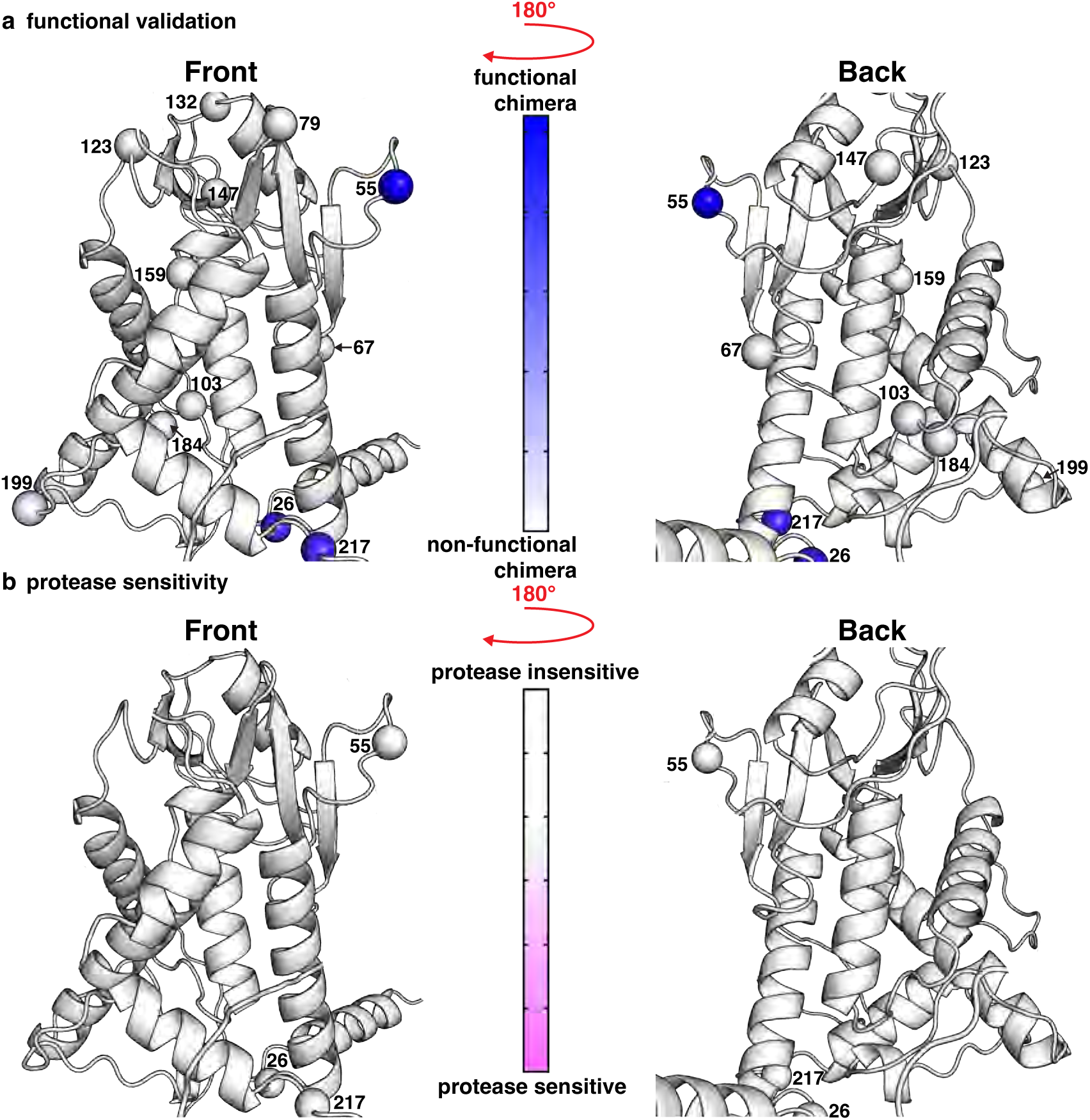
Ste12 HCV Peptide Insertion Sites. **a-b,** Structural views of AlphaFold2 model of Ste12 (AF-P13574-F1-model_v4) with PAM-scan chimeric insertion positions labeled by codon number and represented as spheres. The model is cropped to display only the regions with confident structural predictions.

**Fig. S13.**
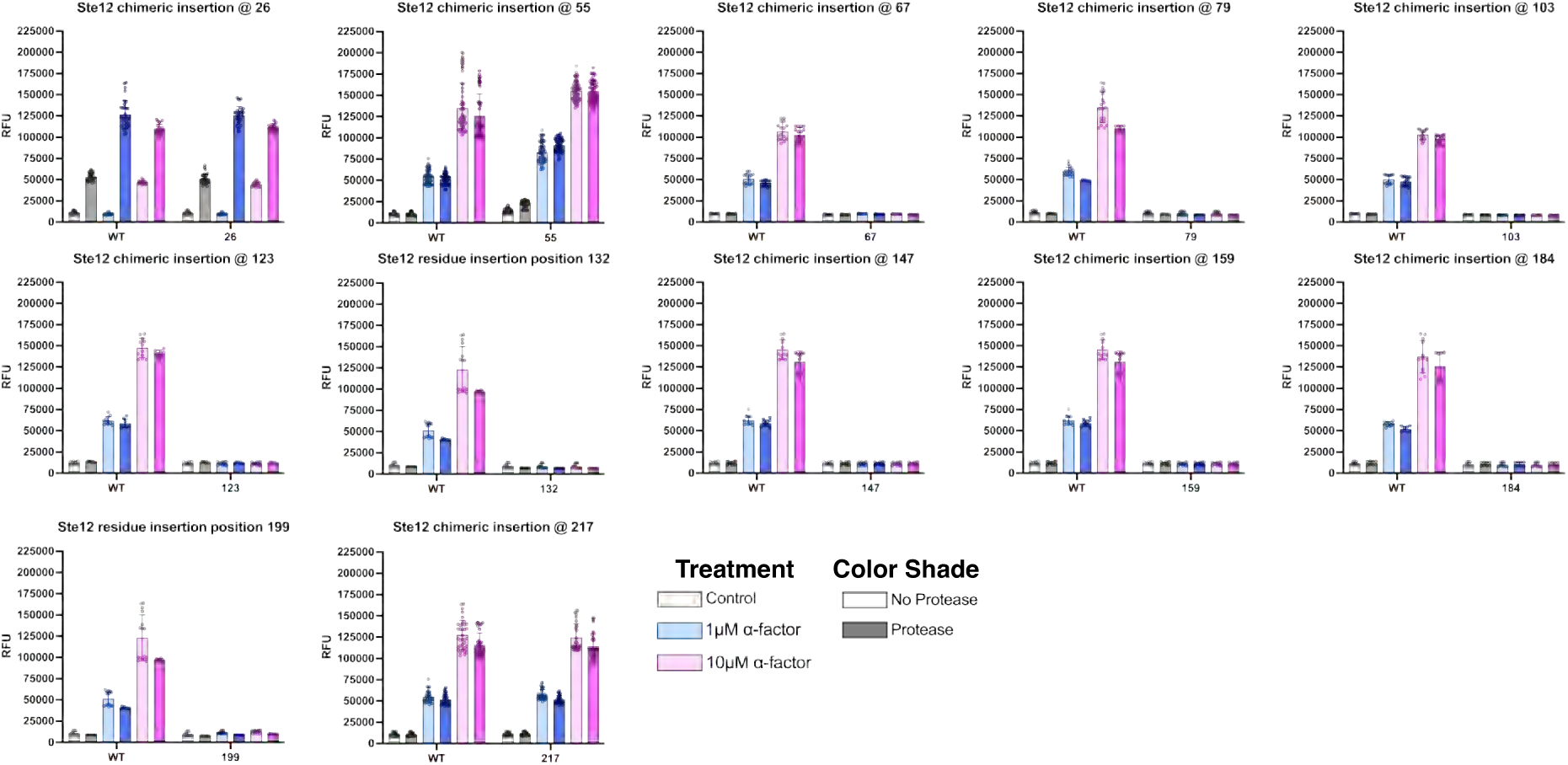
Primary functional validation data for Ste12 HCV peptide chimeras. Pheromone pathway activation was quantified by mTq2 fluorescence and reported as relative fluorescence units (RFU). Each condition includes a minimum of n = 3 independent experiments, each with n = 4 technical replicates.

**Fig. S14.**
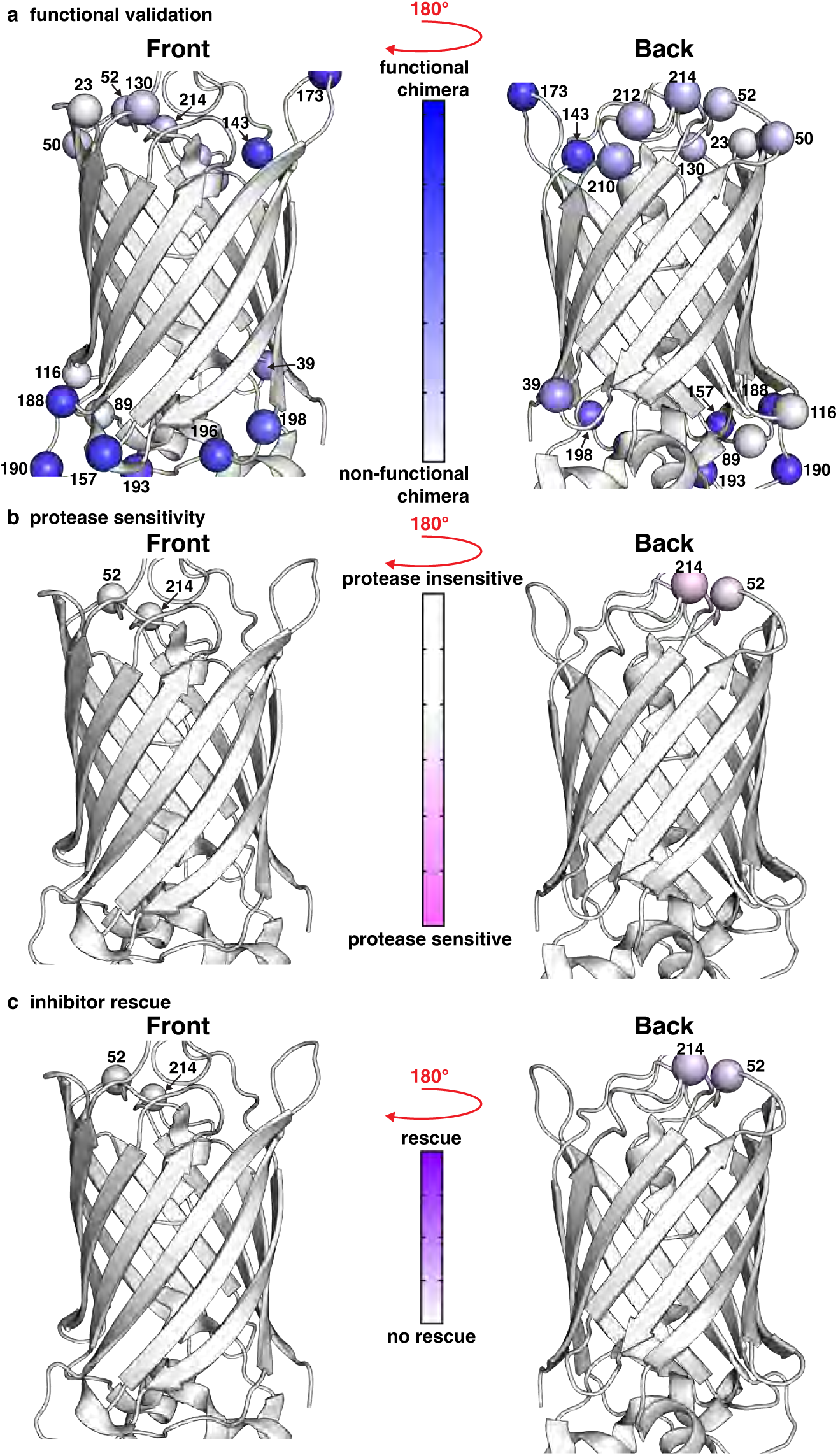
mTq2 HCV Peptide Insertion Sites. **a-b,** Structural views of AlphaFold2 model of mTq2 (PDB: 6yln) with PAM-scan chimeric insertion positions labeled by codon number and represented as spheres.

**Fig. S15.**
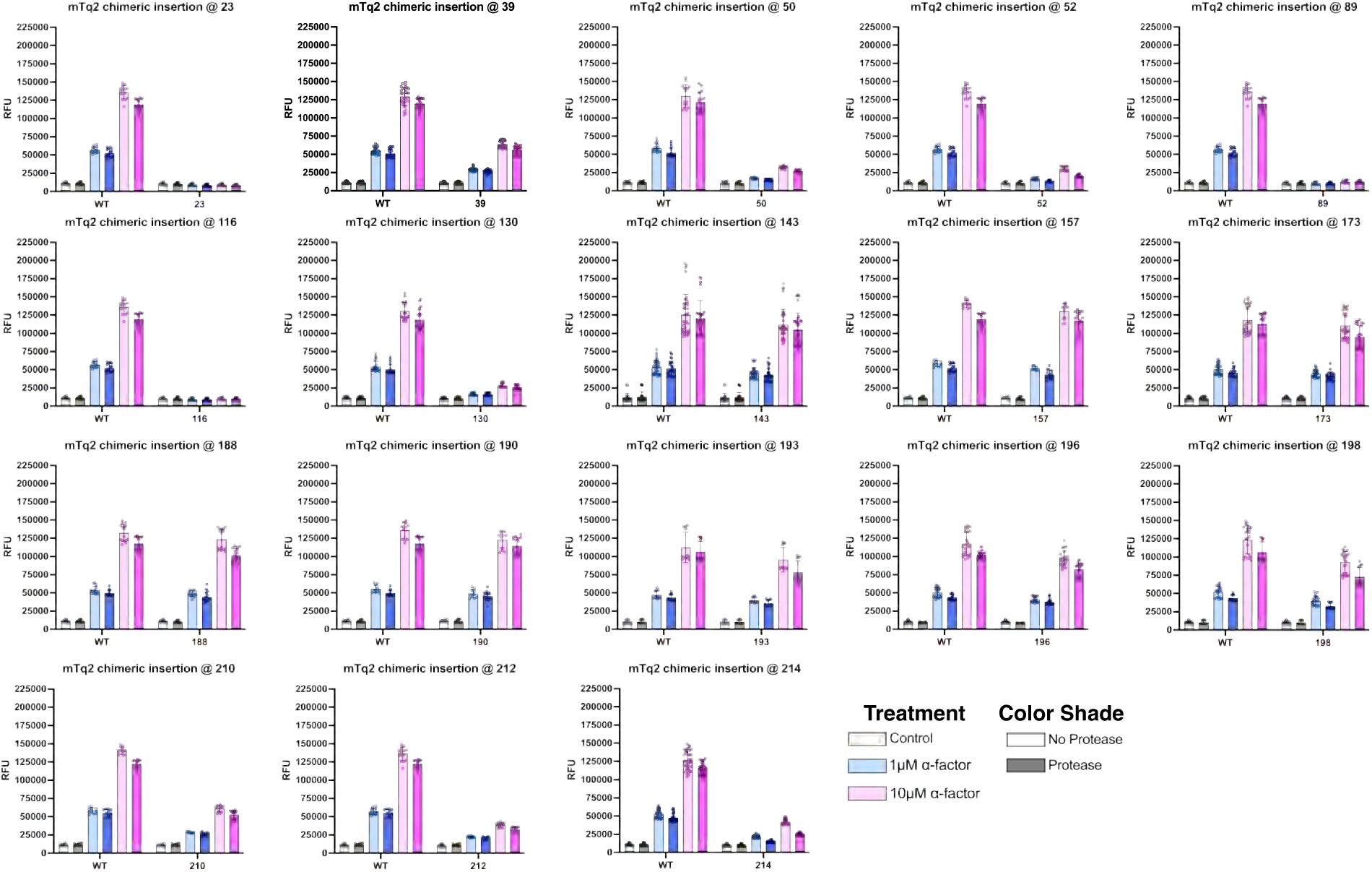
Primary functional validation data for mTq2 HCV peptide chimeras. Pheromone pathway activation was quantified by mTq2 fluorescence and reported as relative fluorescence units (RFU). Each condition includes a minimum of n = 3 independent experiments, each with n = 4 technical replicates.

**Fig. S16.**
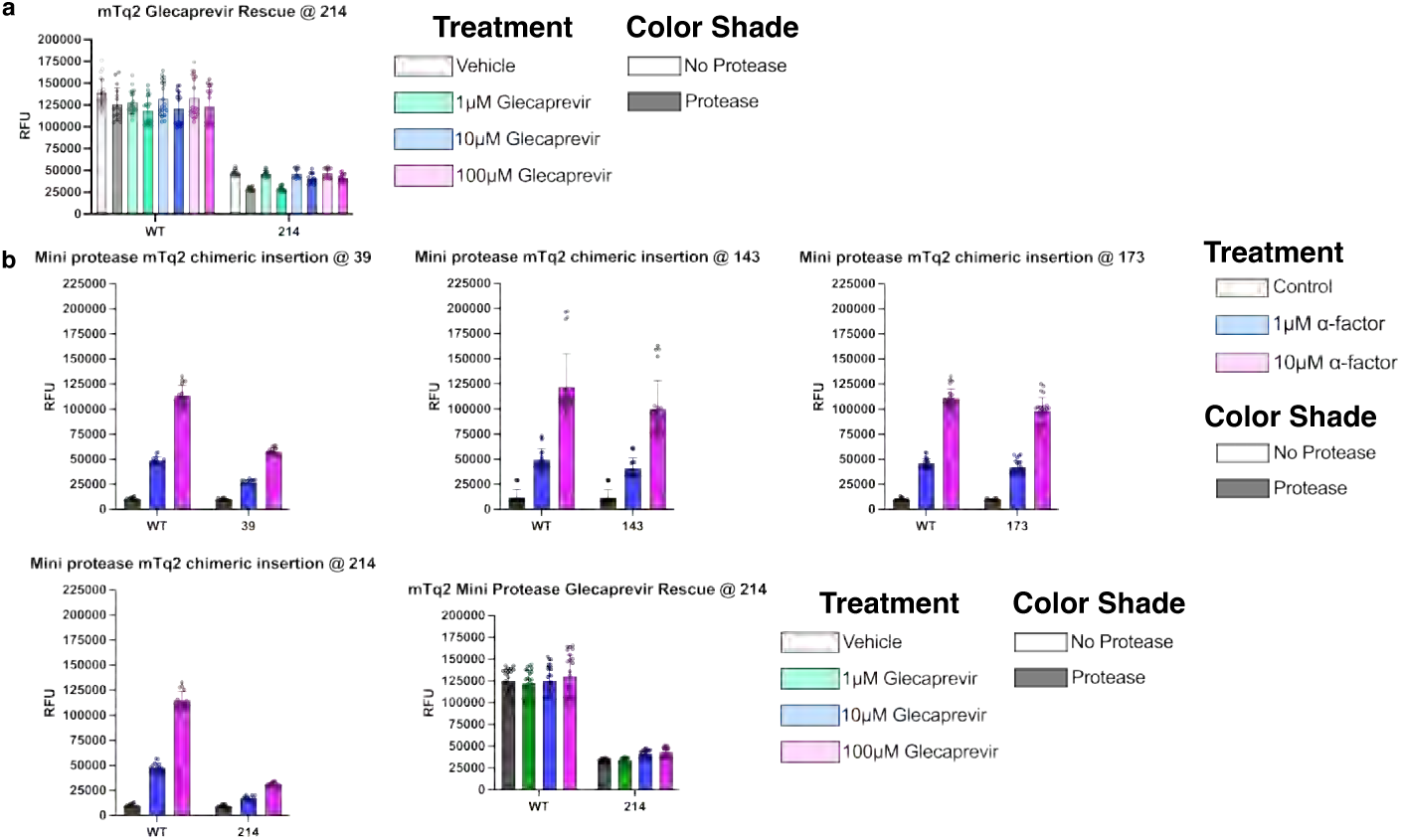
Primary functional validation, protection, and inhibitor rescue data for mTq2 HCV peptide chimeras. **a**, Primary data from the inhibitor assay performed on each mTq2 site that showed reduced protease-dependent signaling. **b**, Primary data from the mini-protease constructs. The α-factor assay (top) was used to derive protease activity, and the inhibitor assay (bottom) was used to calculate rescue by the inhibitor glecaprevir. Pheromone pathway activation was quantified by mTq2 fluorescence and reported as relative fluorescence units (RFU). Each condition includes a minimum of n = 3 independent experiments, each with n = 4 technical replicates.

## References

1 Tang, H. et al. Fusion genes in cancers: Biogenesis, functions, and therapeutic implications. Genes Dis 12, 101536 (2025). 10.1016/j.gendis.2025.101536

2 Chalfie, M., Tu, Y., Euskirchen, G., Ward, W. W. & Prasher, D. C. Green fluorescent protein as a marker for gene expression. Science 263, 802–805 (1994). 10.1126/science.8303295

3 Tsien, R. Y. The green fluorescent protein. Annu Rev Biochem 67, 509–544 (1998). 10.1146/annurev.biochem.67.1.509

4 de Wet, J. R., Wood, K. V., DeLuca, M., Helinski, D. R. & Subramani, S. Firefly luciferase gene: structure and expression in mammalian cells. Mol Cell Biol 7, 725–737 (1987). 10.1128/mcb.7.2.725-737.1987

5 Nakai, J., Ohkura, M. & Imoto, K. A high signal-to-noise Ca(2+) probe composed of a single green fluorescent protein. Nat Biotechnol 19, 137–141 (2001). 10.1038/84397

6 Nikolaev, V. O., Bunemann, M., Hein, L., Hannawacker, A. & Lohse, M. J. Novel single chain cAMP sensors for receptor-induced signal propagation. J Biol Chem 279, 37215–37218 (2004). 10.1074/jbc.C400302200

7 Liu, Y., Witucki, L. A., Shah, K., Bishop, A. C. & Shokat, K. M. Src-Abl tyrosine kinase chimeras: replacement of the adenine binding pocket of c-Abl with v-Src to swap nucleotide and inhibitor specificities. Biochemistry 39, 14400–14408 (2000). 10.1021/bi000437j

8 Irannejad, R. et al. Conformational biosensors reveal GPCR signalling from endosomes. Nature 495, 534–538 (2013). 10.1038/nature12000

9 Wan, Q. et al. Mini G protein probes for active G protein-coupled receptors (GPCRs) in live cells. J Biol Chem 293, 7466–7473 (2018). 10.1074/jbc.RA118.001975

10 DiBerto, J. F., Olsen, R. H. J. & Roth, B. L. TRUPATH: An Open-Source Biosensor Platform for Interrogating the GPCR Transducerome. Methods Mol Biol 2525, 185–195 (2022). 10.1007/978-1-0716-2473-9_13

11 Kroeze, W. K. et al. PRESTO-Tango as an open-source resource for interrogation of the druggable human GPCRome. Nat Struct Mol Biol 22, 362–369 (2015). 10.1038/nsmb.3014

12 Qin, K., Dong, C., Wu, G. & Lambert, N. A. Inactive-state preassembly of G(q)-coupled receptors and G(q) heterotrimers. Nat Chem Biol 7, 740–747 (2011). 10.1038/nchembio.642

13 Masuho, I., Skamangas, N. K., Muntean, B. S. & Martemyanov, K. A. Diversity of the Gbetagamma complexes defines spatial and temporal bias of GPCR signaling. Cell Syst 12, 324–337 e325 (2021). 10.1016/j.cels.2021.02.001

14 Avet, C. et al. Effector membrane translocation biosensors reveal G protein and betaarrestin coupling profiles of 100 therapeutically relevant GPCRs. Elife 11 (2022). 10.7554/eLife.74101

15 Hirabara, S. et al. Clinical efficacy of abatacept, tocilizumab, and etanercept in Japanese rheumatoid arthritis patients with inadequate response to anti-TNF monoclonal antibodies. Clin Rheumatol 33, 1247–1254 (2014). 10.1007/s10067-014-2711-2

16 Gjolberg, T. T. et al. Biophysical differences in IgG1 Fc-based therapeutics relate to their cellular handling, interaction with FcRn and plasma half-life. Commun Biol 5, 832 (2022). 10.1038/s42003-022-03787-x

17 Cohen, M. D. & Keystone, E. Rituximab for Rheumatoid Arthritis. Rheumatol Ther 2, 99–111 (2015). 10.1007/s40744-015-0016-9

18 Mostkowska, A., Rousseau, G. & Raynal, N. J. Repurposing of rituximab biosimilars to treat B cell mediated autoimmune diseases. FASEB J 38, e23536 (2024). 10.1096/fj.202302259RR

19 June, C. H. & Sadelain, M. Chimeric Antigen Receptor Therapy. N Engl J Med 379, 64–73 (2018). 10.1056/NEJMra1706169

20 Lim, W. A. & June, C. H. The Principles of Engineering Immune Cells to Treat Cancer. Cell 168, 724–740 (2017). 10.1016/j.cell.2017.01.016

21 Roybal, K. T. et al. Engineering T Cells with Customized Therapeutic Response Programs Using Synthetic Notch Receptors. Cell 167, 419–432 e416 (2016). 10.1016/j.cell.2016.09.011

22 Jinek, M. et al. A programmable dual-RNA-guided DNA endonuclease in adaptive bacterial immunity. Science 337, 816–821 (2012). 10.1126/science.1225829

23 Cong, L. et al. Multiplex genome engineering using CRISPR/Cas systems. Science 339, 819–823 (2013). 10.1126/science.1231143

24 Mali, P. et al. RNA-guided human genome engineering via Cas9. Science 339, 823–826 (2013). 10.1126/science.1232033

25 Hsu, P. D., Lander, E. S. & Zhang, F. Development and applications of CRISPR-Cas9 for genome engineering. Cell 157, 1262–1278 (2014). 10.1016/j.cell.2014.05.010

26 Kapolka, N. J. et al. Proton-gated coincidence detection is a common feature of GPCR signaling. Proc Natl Acad Sci U S A 118 (2021). 10.1073/pnas.2100171118

27 Kapolka, N. J., Taghon, G. J. & Isom, D. G. Advances in yeast synthetic biology for human G protein-coupled receptor biology and pharmacology. Curr Opin Biotechnol 88, 103176 (2024). 10.1016/j.copbio.2024.103176

28 Kapolka, N. J. et al. DCyFIR: a high-throughput CRISPR platform for multiplexed G protein-coupled receptor profiling and ligand discovery. Proc Natl Acad Sci U S A 117, 13117–13126 (2020). 10.1073/pnas.2000430117

29 Rowe, J. B., Kapolka, N. J., Taghon, G. J., Morgan, W. M. & Isom, D. G. The evolution and mechanism of GPCR proton sensing. J Biol Chem 296, 100167 (2021). 10.1074/jbc.RA120.016352

30 Rowe, J. B., Lee, K. & Isom, D. G. PIONEER: A periplasmic display platform for synthetic biology-based screening of genetically encoded protein regulators. J Biol Chem 302, 110967 (2026). 10.1016/j.jbc.2025.110967

31 Rowe, J. B., Taghon, G. J., Kapolka, N. J., Morgan, W. M. & Isom, D. G. CRISPR-addressable yeast strains with applications in human G protein-coupled receptor profiling and synthetic biology. J Biol Chem 295, 8262–8271 (2020). 10.1074/jbc.RA120.013066

32 Rostamian, H. et al. Rapid CRISPR-Cas9 Genome Editing in S. cerevisiae. bioRxiv (2026). 10.64898/2026.03.27.714888

33 Hvidt, A. & Nielsen, S. O. Hydrogen exchange in proteins. Adv Protein Chem 21, 287–386 (1966). 10.1016/s0065-3233(08)60129-1

34 Krishna, M. M., Hoang, L., Lin, Y. & Englander, S. W. Hydrogen exchange methods to study protein folding. Methods 34, 51–64 (2004). 10.1016/j.ymeth.2004.03.005

35 Li, R. & Woodward, C. The hydrogen exchange core and protein folding. Protein Sci 8, 1571–1590 (1999). 10.1110/ps.8.8.1571

36 Park, C. & Marqusee, S. Pulse proteolysis: a simple method for quantitative determination of protein stability and ligand binding. Nat Methods 2, 207–212 (2005). 10.1038/nmeth740

37 Isom, D. G., Marguet, P. R., Oas, T. G. & Hellinga, H. W. A miniaturized technique for assessing protein thermodynamics and function using fast determination of quantitative cysteine reactivity. Proteins 79, 1034–1047 (2011). 10.1002/prot.22932

38 Isom, D. G., Vardy, E., Oas, T. G. & Hellinga, H. W. Picomole-scale characterization of protein stability and function by quantitative cysteine reactivity. Proc Natl Acad Sci U S A 107, 4908–4913 (2010). 10.1073/pnas.0910421107

39 Martino, S. D., Petri, G. L. & De Rosa, M. Hepatitis C: The Story of a Long Journey through First, Second, and Third Generation NS3/4A Peptidomimetic Inhibitors. What Did We Learn? J Med Chem 67, 885–921 (2024). 10.1021/acs.jmedchem.3c01971

40 Zephyr, J., Kurt Yilmaz, N. & Schiffer, C. A. Viral proteases: Structure, mechanism and inhibition. Enzymes 50, 301–333 (2021). 10.1016/bs.enz.2021.09.004

41 Taremi, S. S. et al. Construction, expression, and characterization of a novel fully activated recombinant single-chain hepatitis C virus protease. Protein Sci 7, 2143–2149 (1998). 10.1002/pro.5560071011

## Methods References

1 Kapolka, N. J. et al. Proton-gated coincidence detection is a common feature of GPCR signaling. Proc Natl Acad Sci U S A 118 (2021). 10.1073/pnas.2100171118

2 Kapolka, N. J. et al. DCyFIR: a high-throughput CRISPR platform for multiplexed G protein-coupled receptor profiling and ligand discovery. Proc Natl Acad Sci U S A 117, 13117–13126 (2020). 10.1073/pnas.2000430117

3 Rowe, J. B., Kapolka, N. J., Taghon, G. J., Morgan, W. M. & Isom, D. G. The evolution and mechanism of GPCR proton sensing. J Biol Chem 296, 100167 (2021). 10.1074/jbc.RA120.016352

4 Rowe, J. B., Taghon, G. J., Kapolka, N. J., Morgan, W. M. & Isom, D. G. CRISPR-addressable yeast strains with applications in human G protein-coupled receptor profiling and synthetic biology. J Biol Chem 295, 8262–8271 (2020). 10.1074/jbc.RA120.013066

5 Taremi, S. S. et al. Construction, expression, and characterization of a novel fully activated recombinant single-chain hepatitis C virus protease. Protein Sci 7, 2143–2149 (1998). 10.1002/pro.5560071011

